# Revisiting the evolutionary history of pigs via de novo mutation rate estimation by deep genome sequencing on a three-generation pedigree

**DOI:** 10.1101/2021.03.29.437103

**Authors:** Mingpeng Zhang, Qiang Yang, Huashui Ai, Lusheng Huang

**Author notes:** The corresponsding authors. State Key Laboratory for Swine Genetic Improvement and Production Technology, Jiangxi Agricultural University, Nanchang, China, 330045.

## Abstract

The mutation rate used in the previous analyses of pig evolution and demographics was cursory and brought potential bias in inferring its history. Herein, we estimated de novo mutation rate of pigs using high-quality whole-genome sequencing data from nine individuals in a three-generation pedigree through stringent filtering and validation. The estimated mutation rate was 3.6 × 10^−9^ per generation, corresponding to 1.2 × 10^−9^ per site per year. Using this mutation rate, we re-investigated the evolutionary history of pigs. Our estimates agreed to the divergence time of ~10 kiloyears ago (Kya) between European wild and domesticated pigs, consistent with the domestication time of European pigs based on archaeological evidence. However, other divergence events inferred here were not as ancient as previously described. Our estimates suggested that: Sus speciation occurred ~1.36 Million years ago (Mya); European pigs split up with Asian ones only ~219 Kya; South and North Chinese wild pig split ~25 Kya. Meanwhile, our results showed that the most recent divergence event between Chinese wild and domesticated pigs occurred in the Hetao plain, North China, approximately 20 Kya, supporting the possibly independent domestication in North China along the middle Yellow River. We also found the maximum effective population size of pigs was ~6 times larger than the previous estimate. Notably by simulation, we confirmed an archaic migration from other Sus species originating ~ 2 Mya to European pigs during pigs’ western colonization, which possibly interfered with the previous demographic inference. Our findings advance the understanding of pig evolutionary history.

## Introduction

*Sus scrofa* (wild boars and domestic pigs) is a subfamily of Suidae, a widespread pig species group of Cetartiodactyla originated in the Oligocene at least 20 million years ago (Mya). Larson et al. (2005), Groenen et al. (2012) and Frantz et al. (2013) made a significant contribution to and systemically illustrated the evolutionary history of pigs. *Sus scrofa* originated on the Island South East Asia (ISEA) during the early Pliocene climatic fluctuations about 3 to 4 Mya (Groenen et al. 2012). The oldest diverging lineage of pigs found to date is of a wild boar population from the North of Sumatra, which split from the Eurasian wild boars around 1.6 to 2.4 Mya (Frantz et al. 2013). Over the past one million years, *Sus scrofa* spread into and colonized almost the entire Eurasian continent (Frantz et al. 2013; Groenen 2016). North and South Chinese *Sus scrofa* populations separated from each other during the Ionian stage approximately 0.6 Mya (Frantz et al. 2013). The domestication of pigs is one of the critical events in the history of human agricultural civilization. Pigs were domesticated in at least two locations: Anatolia (Near East) and China. Pig domestication in Anatolia was well documented, which was indicated at ~10 kiloyears ago (Kya) based on archaeological evidence (Giuffra et al. 2000; Larson et al. 2005; Frantz et al. 2019), while pig domestication in China happened at least 8 Kya based on zooarchaeological analyses from middle China (Jing and Flad 2002; Larson et al. 2007). However, studies on domestication of Chinese wild boars based on genomic analyses were still limited. Nowadays, pigs distribute almost all over the world (Yang et al. 2017). Pigs have lived closely with humans for at least 10,000 years (Larson et al. 2010; Frantz et al. 2013), have been one of the essential providers of animal protein for humans (Huang et al. 2020), and serve as ideal biomedical models for human diseases (Walters et al. 2017).

An accurate mutation rate plays a significant role in understanding many critical questions in evolutionary and population genetics, including effective population size, divergence time, and migration between populations (Lynch 2010a). The two conventional methods used to estimate the mutation rate are: (1) phylogenetic approaches, in which the rate of neutral sequence divergence is equal to the rate of mutation (Kimura 1968); (2) direct detection of the spontaneous germline mutations in a known pedigree (Keightley et al. 2014; Smeds et al. 2016; Pfeifer 2017; Koch et al. 2019), which was used in this study. The latter method, benefiting from the popularity of high-throughput sequencing technologies, has many advantages over the former one (Smeds et al. 2016; Koch et al. 2019). A directed per-generation mutation rate derived from a known pedigree has taken an essential part in effectively revising human history (Scally and Durbin 2012) and dogs (wolves) evolutionary history (Koch et al. 2019).

However, at present, there is still no research specializing in the mutation rate of pigs. The mutation rate used in almost all previous demographics of pigs was set as 2.5 × 10^−8^ per generation same to the default one of humans, and 5 years was used as one generation time of pigs (Groenen et al. 2012; Frantz et al. 2013; Li et al. 2013; Bosse et al. 2014; Nuijten et al. 2016). This resulted in an abnormally sizeable annual mutation rate (5 × 10^−9^) of pigs, which is twice the average value of mammals (Kumar and Subramanian 2002) and is specifically 3.3- and 2.6-fold to that of wolves (Koch et al. 2019) and yaks (Qiu et al. 2015), respectively (fig. 1). The inaccuracy of mutation rate may bring bias in the inference of pig demographic parameters and evolutionary history. This study aims to estimate the mutation rate of pigs directly using the genomes of nine individuals from a three-generation pig pedigree. Based on this mutation rate, we re-investigated the evolutionary and domestication history of pigs through genomic analyses.

**Figure 1.**
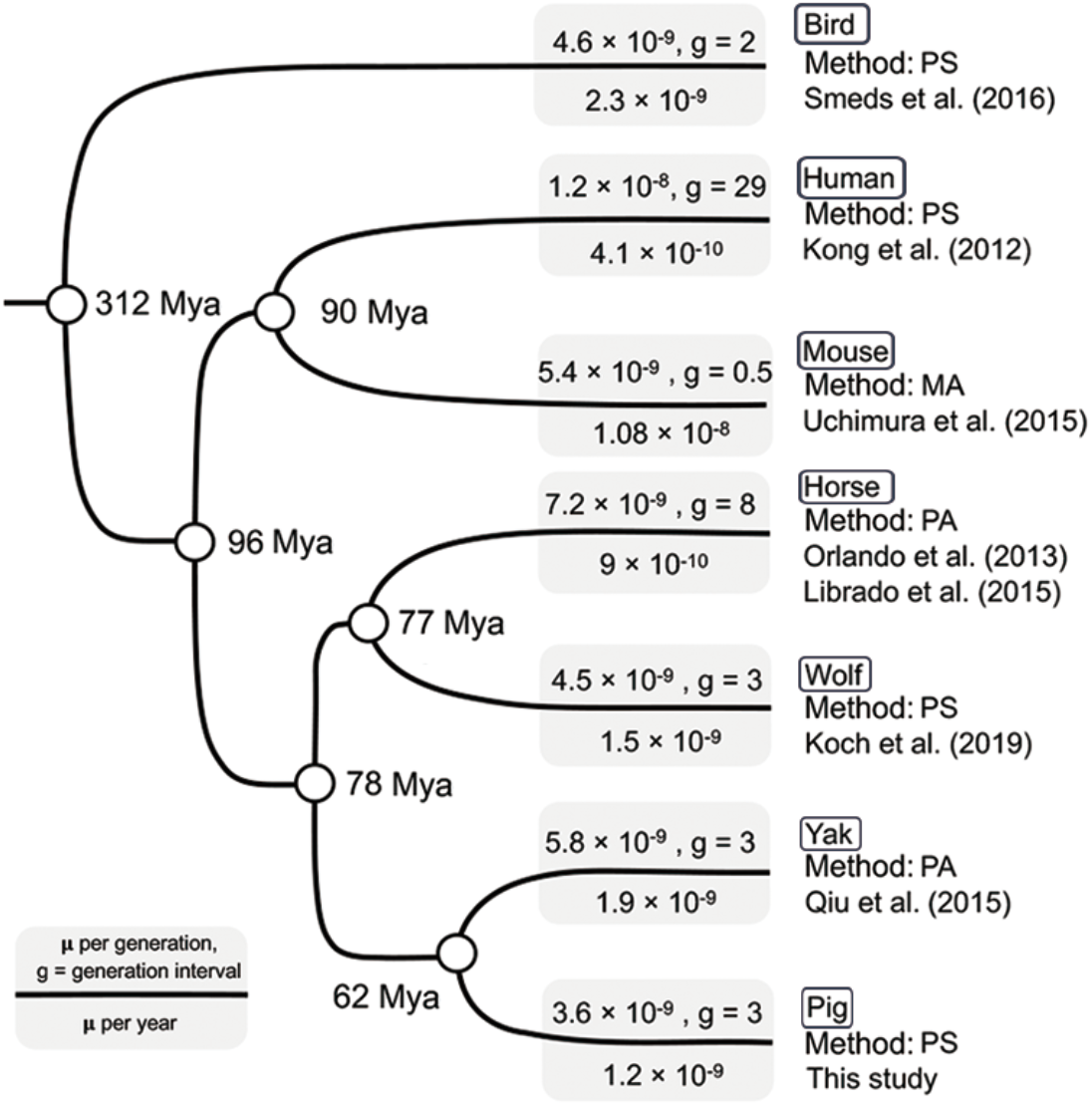
The mutation rate and generation interval used in the demographic inference in birds and several mammals. Different methods were used to estimate mutation rate: PS, pedigree sequencing; MA, sequencing of mutation accumulation lines; PA, Phylogenetic approach.

## Results

### Identification and validation of de novo mutations

A complete three-generation pedigree consisting of nine pigs (4 parents, 2 children, and 3 grandchildren; fig. 2) was re-sequenced with depth more than 20×. In the pedigree, two boars in the parent generation (F0) were White Duroc, and two F0 sows were Erhualian. We applied highly stringent filtering criteria as previously described (Smeds et al. 2016) to carefully screen for de novo mutations (DNMs). In total, 44 DNMs (supplementary table S1 and S2, Supplementary Material online) were identified in the child (F1) and grandchild (F2) generation pigs (7 to 11 DNMs per individual), which were homozygous for the reference allele in all F0 individuals. None of these DNM sites were known to segregate when searching in the *Sus scrofa* dbSNPs 150.

**Figure 2.**
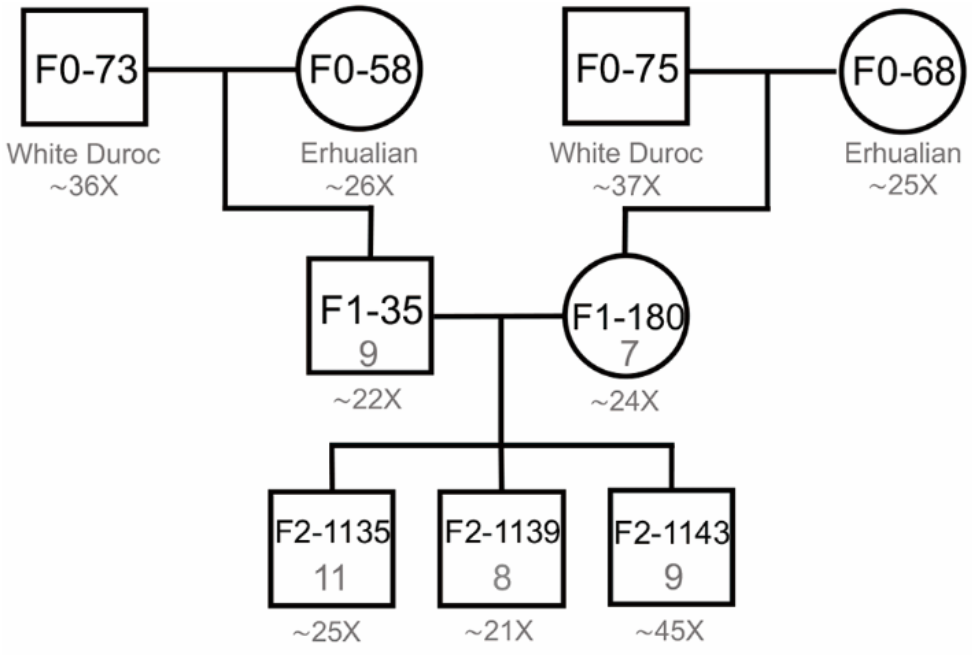
Whole-genome sequences from nine pigs of known pedigree were analyzed to detect de novo mutations. The characters and numbers in the square (male) and circle (female) indicate pig ID and the number of de novo mutations, respectively. Below each individual is its breed and the average sequencing depth of coverage (see Supplementary information, Table S3, for more detail of per individual).

In the F1 generation, 7 and 9 variants in F1-180 and F1-35, respectively, passed manual curation (supplementary table S1, Supplementary Material online). F1-180 transmitted all 7 mutations to the F2 offspring, while one mutant out of 9 in F1-35 was not transmitted to any of 3 offspring (supplementary table S1, Supplementary Material online). We detected a total of 28 mutants passed all the bioinformatic filtering criteria in the F2 generation and these mutants also passed manual curation (fig. 2; supplementary table S1 and S2, Supplementary Material online).

We applied Sanger sequencing to check the de novo mutations further. 40 out of the 44 mutants were validated by Sanger sequencing, including the mutant in F1 that was not transmitted to any of 3 offspring (supplementary table S1 and S2, Supplementary Material online). The remaining 4 mutants (1, 1, and 2 in F1-35, F2-1135 and F2-1139, respectively) were invalidated and detected as homozygotes for the reference allele by Sanger sequencing. In the mapping results of resequencing data, the ratios of the mapped reads supporting alternative allele to all the reads at these 4 sites were 6/17, 10/30, 4/14 and 3/11, respectively (supplementary fig. S1, S2, S3 and S4, Supplementary Material online). Among them, the mutant at chr7:46553490 was even supported by stable inheritance in the F2 generation (supplementary fig. S1, Supplementary Material online). We found the mutant at chr3:9293354 with the ratio of 3/11, same to the ratio value of the invalidated mutant at chr15:107528000, was detected as real (supplementary table S2, Supplementary Material online). Thus, we did not exclude these four mutants. One possible explanation might be the bias in the sequencing results caused by PCR errors before Sanger sequencing (Ikegawa et al. 2002).

Furthermore, we explored the characteristics of the 44 DNMs. There were 14 mutations in intergenic regions, 25 in introns, one in 3’-UTR regions, one in splicing regions and three in the coding sequence. Among the three exonic sites (supplementary table S1, Supplementary Material online), one mutation was non-synonymous in Piccolo Presynaptic Cytomatrix Protein (PCLO), a part of the presynaptic cytoskeletal matrix. There were 13 A:T>G:C and 28 G:C>A:T mutations (supplementary table S1, Supplementary Material online), confirming a mutation pressure in the direction of A+T previously seen in both eukaryotes (Lynch 2010b; Smeds et al. 2016) and prokaryotes (Hershberg and Petrov 2010).

### The mutation rate in pigs

A total of 44 DNMs were observed in 10 transmissions, with an average of 4.4 mutations per transmission. Among the five offspring, the effective sequences for screening after filtering ranged between 1.17-1.28 Gb, with an average size of 1.23 Gb, approximately representing 54.6% of pig autosomal genome (see details in Methods; supplementary table S3, Supplementary Material online). The parts containing repetitive and not meeting filter criteria for coverage and quality were excluded. Finally, the mutation rate was calculated to be 3.6 × 10^−9^ per base pair per generation.

Pigs are typically social animals, living in the form of polygamy (Canu et al. 2015). The ages of estrus in sows and boars in the wild are different. Age at first pregnancy varies in the wild from about 10-20 months (Singer 1981), while boars begin rut when they are 3-5 years. The first rut age of 4-5 years was documented in Russian wild boars by Heptner et al. (1988) and 3-4 years was recorded in Chinese wild boars (Wang 2012). Pigs are multiparous animals, and the gestation period lasts about 114-130 days (Comer and Mayer 2009). Comparing to the animals with single birth, like cattle and yak, we think the generation transmission of pigs is of good continuity. Therefore, we set the generation interval of pigs as 3 years, which is roughly equivalent to the average age of the first pregnancy in sows and the beginning rut in boars plus the pregnant gestation period of sows. We noticed that evolutionary studies of dogs (wolves) and yak, which are in a close phylogenic distance with pigs, also adopted 3 years as their generation interval (Freedman et al. 2014; Qiu et al. 2015), suggesting the reasonableness of 3 years as generation interval of pigs. Based on this generation interval, we obtained an annual mutation rate of 1.2 × 10^−9^, close to the mutation rate (1.5 × 10^−9^; the mutation rate was estimated via a known pedigree) of wolves (Koch et al. 2019). The annual mutation rate of pigs is in the same order of magnitude as those of mammals (fig. 1) and lower than the mean mutation rate (2.2 × 10^−9^) of mammals (Kumar and Subramanian 2002).

Additionally, we employed the phylogenetic approach to estimate the mutation rate of pigs, following the procedure of mutation rate estimation used in dogs (Wang et al. 2016). The mutation rate of pigs was estimated to be 1.53 × 10^−9^ per year (supplementary table S4, Supplementary Material online). In humans, the mutation rate obtained from whole-genome pedigree data is lower than those obtained from the phylogenetic approaches (Nachman and Crowell 2000), coinciding with the results in dogs and wolves (Wang et al. 2016; Koch et al. 2019). Here we obtained a slightly lower mutation rate from whole-genome pedigree data than the estimate using the phylogenetic approach, which is in line with those previous studies in humans and dogs and also suggesting the high accuracy of the de novo mutation rate in pigs.

### Demography of Sus species

We took the de novo mutation rate as a parameter in the pairwise sequentially Markovian coalescent model (PSMC) (Li and Durbin 2011) and the multiple sequential Markovian coalescent model (MSMC) (Schiffels and Durbin 2014) to reconstruct the population history of Sus (see details in Methods). Sumatran wild boar (SMW), European wild boar (EUW), North Chinese wild boar (NCW) and South Chinese wild boar (SCW) represented *Sus scrofa* of different geographic distributions in this study. *Sus cebifrons*, as an outgroup, was involved to date the speciation of *Sus scrofa* from Sus. Each breed contained two individuals (supplementary table S5, Supplementary Material online).

PSMC exhibited messy-looking demographic trajectories: demographic trajectories of different pigs began to separate at 2 Mya, except for that of North and South China pigs, which began to separate from each other ~200 Kya (fig. 3A). Such messy curves possibly suggested a short common history among different breeds of pigs, and indicated they diverged from each other million years ago. However, we could find common points like all pigs peaked with effective population size (Ne) during 1 ~ 2 Mya. *Sus cebifrons*, SMW and EUW have a similar Ne of ~ 2.7 × 10^5^ during the peak period, while Chinese pigs have a lower Ne then. The maximum estimation of effective population size here is ~6 times larger than that estimated before (Groenen et al. 2012; Frantz et al. 2013). Thereafter, *Sus cebifrons* and SMW experienced a rapid population decline. EUW and SCW experienced a lighter decline and then stayed population stability or rising beginning 400 Kya. Notably, the trajectory of EUW was similar in trends with that of Chinese pigs before ~ 300 Kya but showed relatively higher Ne. All pigs suffered a bottleneck during the Last Glacial Maximum (LGM; 20 Kya; fig. 3A).

**Figure 3.**
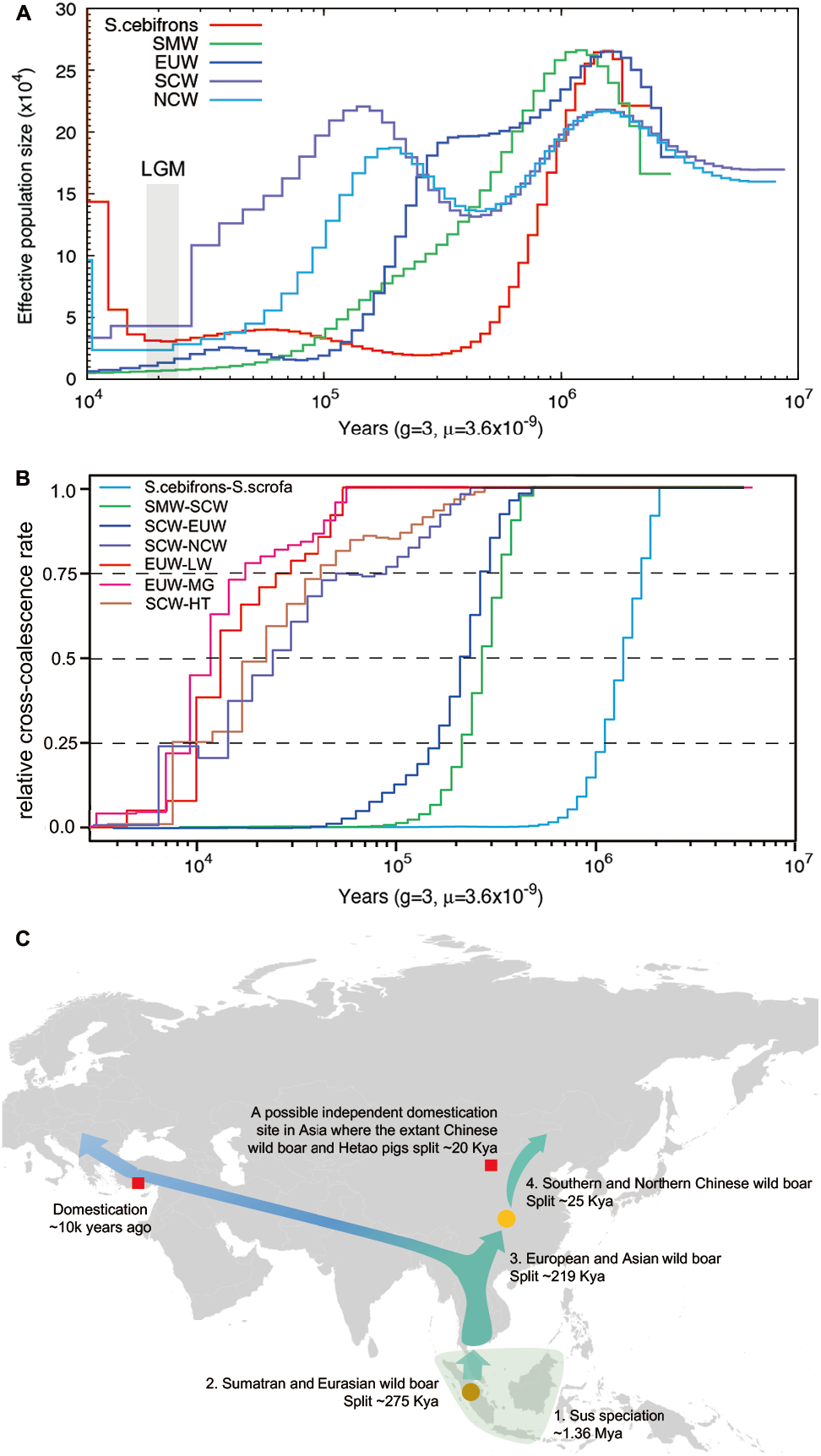
Population history of Sus species. (A) show the changes in effective population size of *Sus cebifrons* and the wild pigs over past year inferred by PSMC. The Last Glacial Maximum (LGM) is highlighted in grey. (B) Split time for population pairs estimated by MSMC. A relative cross-coalescence rate of 0.5 is defined as the divergence time. The cold color lines indicated the split time of wild pig breeds in different regions; the warm color lines could somehow reflect the domestication time, even though the extant wild pigs may not be the domesticated breeds’ direct ancestors. See Supplementary information, Table S6, for more detail of per breed. (C) A map depicting the hypothetical spread of wild pigs across Eurasia and the domestication events of pigs happened in the Middle East and China. The shade covered in Southeast Asia indicates Sus originated here. The circles represent the “node” groups, connecting two different groups. The brown circle depicts Sumatran wild pigs. The yellow circle depicts South Chinese wild pigs. The red squares refer to the domestication events.

MSMC let us study the genetic separation between two populations as a function of time based on relative cross-coalescent rates (RCCR). The RCCR curve reached a value of 0.5 at ~ 1.36 Mya for the comparison of *Sus cebifrons* and *Sus scrofa* (SCW; fig. 3B; supplementary table S6, Supplementary Material online), indicating Sus speciation occurred during this period on ISEA. According to the 0.5 RCCR cutoff defined as the divergence time, we could also judge that the divergence time between SMW and SCW was ~ 275 Kya, EUW and Asian wild boar (ASW) separated ~ 219 Kya, NCW and SCW split ~ 25 Kya, and European wild and domesticated pig diverged ~ 1.3 Kya (fig. 3B; supplementary table S5, Supplementary Material online).

We noticed that the divergence time indicated by MSMC was largely more recent than that implied by trajectories of PSMC. To further explore this issue, we took the comparison of EUW and SCW as an example. Although the results of PSMC showed that the demographic curves of EUW and SCW separated ~ 2 Mya (Supplementary fig. S5A, Supplementary Material online), RCCR in MSMC hasn’t reached 0.5 until ~ 219 Kya, indicating that Eurasian pigs did not really split ~ 2 Mya, but split 219 Kya (supplementary fig. S5B, Supplementary Material online). We applied MSMC-IM software, fitting a continuous model to coalescence rates to estimate gene flow within and across pairs populations, to confirm the time of divergence event. It still showed a peak at ~ 200 Kya rather than 2 Mya, indicating Eurasian pigs’ split time was ~ 200 Kya (supplementary fig. S5C, Supplementary Material online).

We estimated the divergence time between wild boar and domesticated pig in Europe and China, respectively. We used EUW (Netherlands wild pigs) paired with two different domesticated pig breeds (Large White and Mangalica pigs), which are located at the most proximal and distal branch relative to the cluster of European wild pigs, respectively (supplementary fig. S6, Supplementary Material online). Their divergence times were estimated to be 13,482 and 11,204 years ago, respectively, coinciding with the generally accepted domestication time of ~10 Kya (Groenen et al. 2012; Frantz et al. 2013; Frantz et al. 2019). However, the direct ancestors of European domestic pigs are not the existing European wild boars, but the extinct wild boars from the Middle East (Larson et al. 2005). The divergence time is expected to be older than the domestication period of 10 Kya, in line with the estimated date of Large White and Mangalica pigs splitting from European wild boars. We also tested divergence time between the different geographical distributed Chinese domesticated pig breeds and Chinese wild pigs, including ones from North China and South China (supplementary fig. S7 and table S5, Supplementary Material online). We found the domesticated pig breed on Hetao plain, located at the intersection of the middle Yellow River and Inner Mongolia, most recently split up with North and South Chinese wild pigs at around 20 Kya (fig. 3B, supplementary fig. S7 and table S6, Supplementary Material online). Interestingly, several studies addressed the possibility of domestication along the middle Yellow River ~8 Kya based on the ancient mitochondrial DNA (Xiang et al. 2017) and archaeological evidence (Larson et al. 2010). Our results further confirmed North China along the middle Yellow River could be a domestication site in Asia. But here we cannot decide domestication time according to the divergence time for the same reason as in European pigs that the ancestor of domesticated pigs might not be the extant wild boars. We also detect a severe bottleneck during the LGM in all domesticated pigs (supplementary fig. S8, Supplementary Material online).

To sum up, we can summarize the evolutionary history of Sus (fig. 3C): Sus speciation occurred on ISEA ~ 1.36 Mya, leading to the emergence of the oldest *Sus scrofa*; then pigs arrived in Eurasia from ISEA and colonized Southeast Asia at ~ 275 Kya; the spread of wild boars into Europe was ~ 219 Kya; the pigs in South China didn’t migrate to North China until ~ 25 Kya. Additionally, North China along the middle Yellow River could be an independent domestication site in Asia where the wild pigs and domesticated pigs split ~20 Kya. The divergence time between European wild and domesticated pig was first estimated at around 10 Kya using genomic data.

### Contradictions in previous evolutionary history of pigs

Frantz et al. (2013) applied an approximate likelihood method as implemented in MCMCtree to estimate divergence time between Sus species, in which they set the splitting time between *Phacochoerus africanusa* and Sus as a root age at 10.5 Mya based on phylogenetic research on mitochondrial DNA of extant sub-Saharan African suids (Gongora et al. 2011). This ancient root age was used to adjust the prior of the mutation rate, which was set to obey a gamma distribution as G(1,125), in Bayesian clock dating. Their divergence time estimates suggested that populations of *Sus scrofa* from Asia migrated west approximately 1.2 Mya. Groenen et al. (2012) and Frantz et al. (2013) also used the result of PSMC (Li and Durbin 2011), in which the population sizes of European and Asian lineages started to diverge around ~ 1 Mya (supplementary fig. S5D, Supplementary Material online), as a vital supporting for the distinct Asian and European pig lineages splitting ~ 1.2 Mya and illustrated an increase in the European population after pigs arrived from Asia.

In this study, we repeated PSMC analyses on Eurasian wild pigs and further applied MSMC software (Schiffels and Durbin 2014) and MSMC-IM (Wang et al. 2020) to make forward and backward arguments for the above studies. First, we used the divergence time between European and Chinese wild pigs (1.2 Mya) estimated by Frantz et al. (2013) to infer the mutation rate for pigs. This resulted in a mutation rate estimation of ~ 6.5 × 10^−10^ per site per generation (fig. 4A), which is an order of magnitude less than the mutation rate of other mammals (fig. 1). We also tried to use this mutation rate to estimate divergence time between European wild boars and domesticated pigs (European wild pigs and Large White). This resulted in a more ancient divergence at ~70 Kya than the domestication time around 10 Kya indicated by archaeological evidence. We noticed the mutation rate applied in PMSC by Groenen et al. (2012) was 2.5 × 10^−8^ per site per generation. This means that two significantly different mutation rates (fig. 4B) reflect a similar divergence history of Eurasian pigs. All of these possibly suggested there were some biases in the previously estimated pig history.

**Figure 4.**
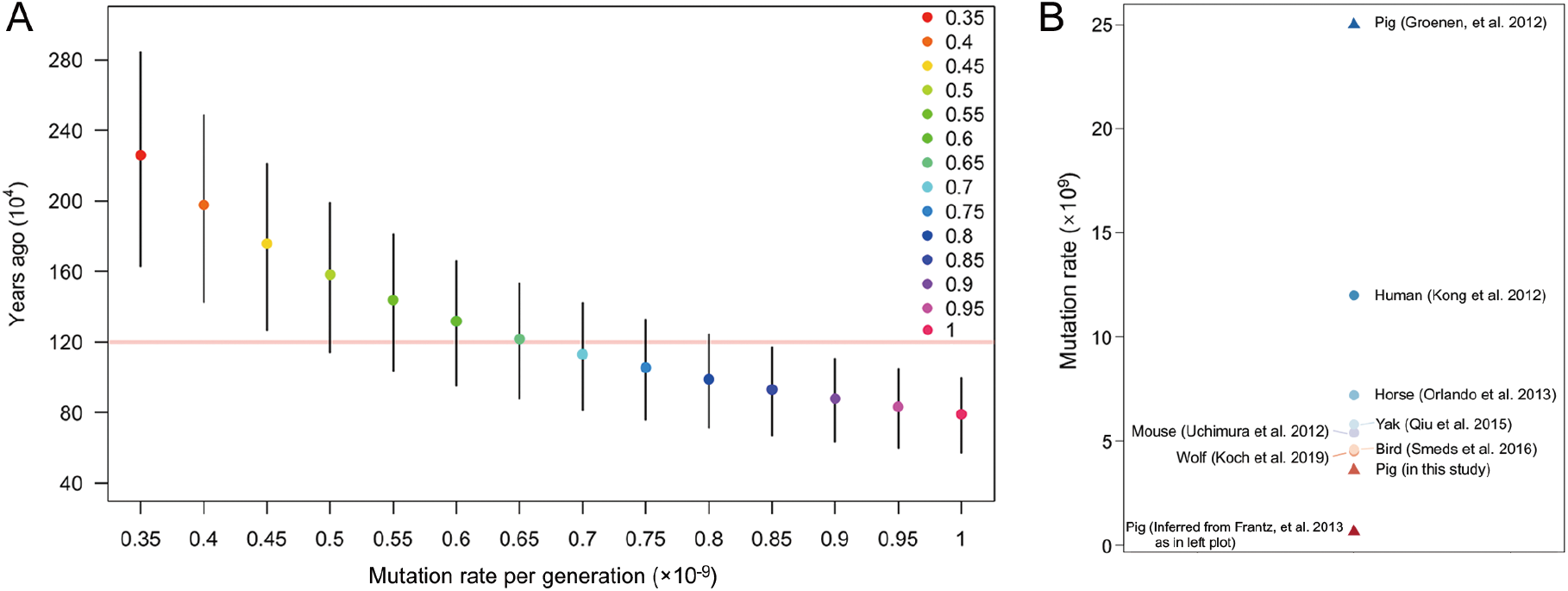
The abnormal mutation rate in the past population studies of pigs. (A) The Pig mutation rate inferred using MSMC when fixing the divergence time between SCW and EUW at 1.2 Million years ago. SCW - EUW divergence time inferred by MSMC using various mutation rates with 3 years fixed as generation interval. Dots, lower and upper bar represents the time at which cross-coalescence rate dropped below, 50%, 25% and 75%, respectively. The red horizontal line represents the SCW - EUW divergence time from Frantz et al. (2013). (B) Scatter plot shows the mutation rate of pigs (triangles) estimated in our study and used in the past research, and exhibits the mutation rate of other species (circles) in Figure 1. Mutation rate here is in a unit of per site per generation.

Then, we run the PSMC, MSMC and MSMC-IM with the mutation rate set as 2.5 × 10^−8^ per generation. In order to compare with the previous results, we only used EUW and SCW here. Surprisingly, we identified four possible contradictions between the previous estimation and the results of MSMC and MSMC-IM here: firstly, both results of MSMC and MSMC-IM indicated a totally different divergence time between European and Chinese wild pigs as before discussed (Groenen et al. 2012): the event of divergence appears at the first cross-over of lines corresponding to the time of ~ 40 Kya (see supplementary fig. S5D, E, and F, Supplementary Material online), rather than the corresponding intersection point at approximately one million years; second, if the event of divergence actually happened at the first cross-over, the date is only 40 Kya, an illogical split time compared to the declared 1 Mya (Groenen et al. 2012; Frantz et al. 2013); third, the divergence time of European wild and domesticated pigs was estimated to only ~ 2 Kya, an unreasonable recent date compared to the domestication time around 10 Kya, via MSMC and MSMC-IM. Last but not least, compared to other mammals, the effective population size of pigs (the maximum Ne was ~ 4 × 10^4^; see supplementary fig. S5D, Supplementary Material online) was largely lower than those of dogs (the maximum Ne of ~ 15 × 10^4^) (Wang et al. 2020) and yak (the maximum Ne of ~ 16 × 10^4^) (Qiu et al. 2015). The ratio of non-synonymous to synonymous heterozygosity (π_N_/π_S_), as a measure of the mutation load, can be applied to the Ne comparisons among distantly related species and was found to be negatively correlated to population size (Galtier and Rousselle 2020). The π_N_/π_S_ ratio of 0.62 - 0.80 in pigs was found lower than that of 1.04 - 1.19 in dogs (Takashi et al. 2018), which meant that pigs had a larger Ne when compared to dogs. This is also contrary to the small Ne estimation of pigs using previous commonly used mutation rate of 2.5 × 10^−8^ per site per generation. We thought the reason why these contradictions occurred could be the use of abnormal large mutation rate in the previous studies.

### Validation of archaic admixture in European *Sus scrofa* by simulations

When we applied the mutation rate of 3.6 × 10^−9^ per site per generation to infer pig demographic history, the MSMC-IM results approximately displayed two pulses (supplementary fig. S5C, Supplementary Material online): one pulse corresponding to the place where RCCR was equal to 0.5, and the other pulse appearing between 1 and 4 Mya, which indicated a migration into SCW or EUW from an archaic population. Correspondingly, recent evidence showed pygmy hogs and a now-extinct Sus species interbred with *Sus scrofa*, suggesting that inter-species admixture accompanied the rapid spread of wild boars across mainland Eurasia and North Africa (Liu et al. 2019). Thus, wild boar had greater chances of encountering and temporal co-existing with local species during the expansion to Europe, enabling possible inter-species hybridization. The phenomenon that the Ne of EUW at the peak of the PMSC curve is similar to those of pigs on ISEA (fig. 3A) possibly further suggested a migration from an archaic pig or another Sus species to European pigs, instead of to SCW, during the colonization across Eurasian mainland. We suspected that this introgression from the archaic population possibly contributed to the difference in the curve of Ne between EUW and SCW before the point of their separation. To test this hypothesis, we performed a series of simulations in which two populations separated 219 kya,with one receiving a varying level of gene flow from an archaic population (fig. 5). We assumed the third population diverged from the ancestor of the former two populations 500 Kya (fig. 5A) and 2 Mya (fig. 5C), respectively, to check how the different archaic donors could affect the trajectories of Ne of receptor. With simulations under this *split-with-archaic-admixture* model (fig. 5), we found archaic admixture did lead to the uplift of Ne of a period when varing different extent of migration (fig. 5B and D). Specifically, the receptor with archaic admixture from the donor that separates from the ancestral population at 2 Mya (fig. 5D), instead of 500 Kya (fig. 5B) in our simulations, exhibited a similar trajectory with that of EUW. Thus, it is most likely to be another Sus species, diverging ~2 Mya, introgressed into EUW. The uplift of Ne curve attributed to the admixture from an archaic population explained the variation trend of the curves of the EUW and SCW is almost same, but Ne of EUW before the divergence happened at the first cross-over of the curves in PSMC is relatively larger (supplementary fig. S5A and D, Supplementary Material online). This uplift of the regional Ne curve of EUW gave us the illusion of deep divergence between EUW and ASW, which possibly brought the bias on previous inference of pig evolutionary history.

**Figure 5.**
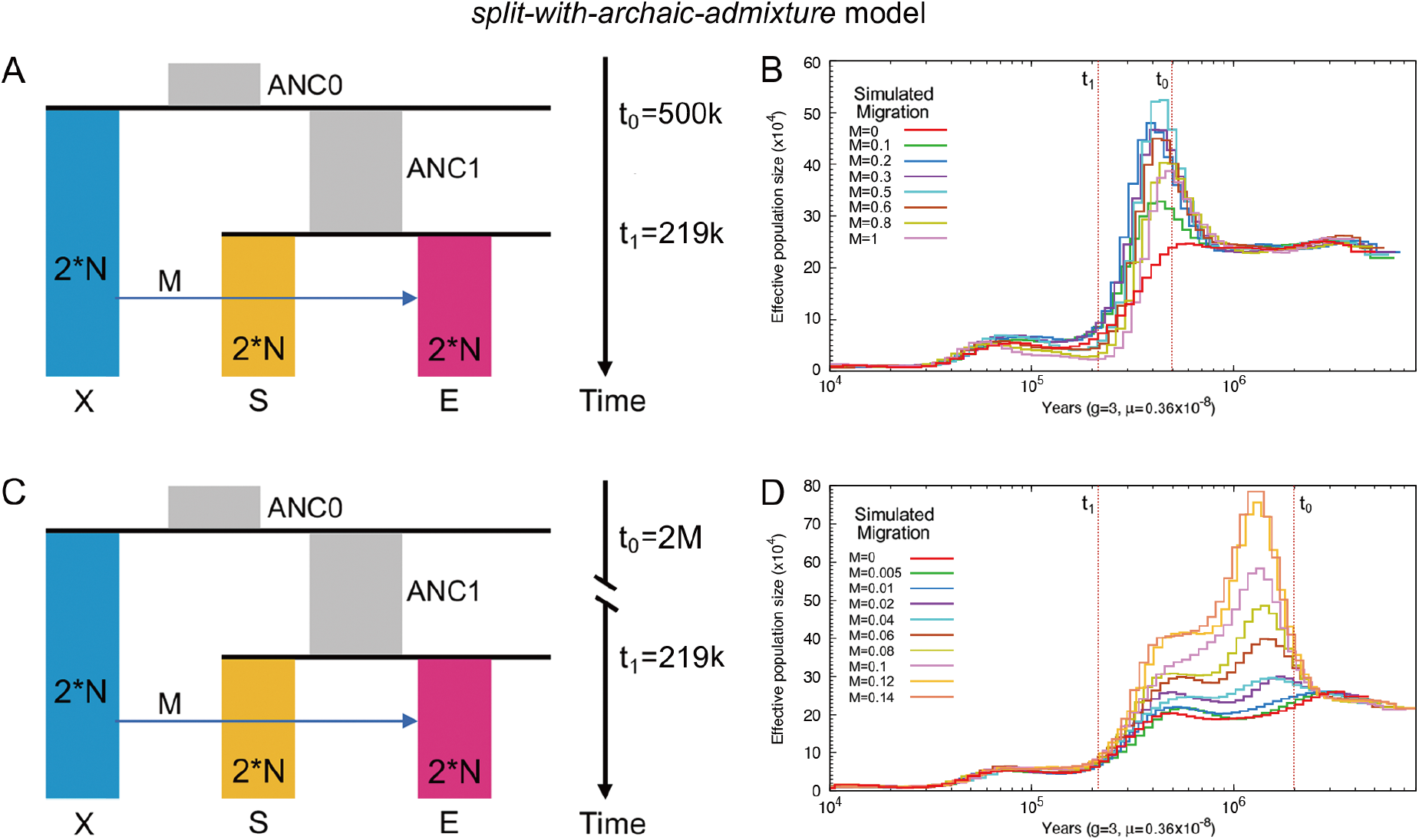
A *split-with-archaic-admixtu*re Model used for simulation testing of whether migration from an ancient population X affect PSMC curves. We simulated data under two scenes with different split time of ANC1 from population X: 500 Kya (A and B) and 2 Mya (C and D). M was migration strength, which was computed in form of 4N*m, and we set M to be different values. PSMC were performed using simulated data of population E. N is the initial population size used to scale parameters in simulation, which is set to 10,000 (default by ms program).

## Discussion

### The accuracy of DNMs detected in a three-generation pedigree by high-quality resequencing

Previous studies have shown that the bioinformatic pipeline we refer to can guarantee the high accuracy of DNMs (Keightley et al. 2014; Keightley et al. 2015; Smeds et al. 2016; Pfeifer 2017). All candidate mutations passed a stringent two-step manual curation by the Integrated Genomics Viewer (IGV) (Robinson et al. 2011). Several procedures were applied to validate mutations during the DNMs identification, including following the stable inheritance of new mutations to subsequent generations and Sanger sequencing. Pfeifer (2017) validated the manually curated candidate mutations in the F1 generation using their stable Mendelian inheritance to the next generation (F2) to exclude the false-positive mutations, even though there was only one individual in F2 of that pedigree. This study had 3 grandchildren with high depths of coverage (21× - 45×) to exclude the false-positive mutations. Additionally, we used another individual F1-43 (with coverage of 40×), only sharing the same father (F0-73) with F1-35, to check the independent genotyping status of the other two individuals (F1-35 and F1-180) in F1, following the criteria that no other individuals in the same generation are heterozygous or homozygous for the alternative allele. We could judge that the false-positive rate of two individuals in the F1 generation was very low based on the following reasons: (1) the use of a stringent bioinformatic pipeline; (2) the final manual inspection by a two-step approach to avoid the interference of the insertion and deletion around candidate mutant; (3) validation by independent genotyping in the same or the previous generation(s); (4) the fact that sites of mutation events were monomorphic in large population samples (dbSNPs 150); (5) the fact that stable inheritance was confirmed for mutants appearing in the F2 generation. Notably, there is one mutant in F1 that was not transmitted to any of the 3 offspring. We detected a total of 28 mutants passed the manual curation in three F2 generation individuals. The proportions of mutations identified in the F1 (16/44 = 36.4%) and F2 generations (63.6%) were approximately in agreement with the proportions of meiosis scored in the F0 (4/10 = 40%) and F1 (6/10 = 60%) generations (supplementary table S1, Supplementary Material online). The slightly higher ratio (63.6%) in the F2 than expected could be attributed to all male individuals, while there is one female, containing 7 mutants, the lowest number of mutants among all the individuals, in the F1 generation. The mutation rate was reported a pronounced male bias in humans and chimpanzee (Kong et al. 2012; Francioli et al. 2015), which possibly explained the slightly lower ratio (36.4%) in F1. Wang and Zhu (2014) also adopt a similar pedigree design to solve the false-positive problem. In this study, we totally identified 44 de novo mutants following the pipeline, of which 40 mutants were validated by Sanger sequencing. Even though validation of four mutants failed by Sanger sequencing, we determined to keep those four ones for the subsequent analysis after balancing the possible unknown errors in sanger sequencing (e.g. PCR errors) and the low false-positive rate following the pipeline. Keightley et al. (2014) addressed the rate of false negatives by adding synthetic mutations to read data from a *D. melanogaster* pedigree containing14 individuals when using a very similar bioinformatic pipeline for mutation identification as in our study. They detected 99.4% of all callable synthetic mutations following the bioinformatic pipeline, suggesting that the rate of false negatives was negligible. Smeds et al. (2016) considered the filtering of both heterozygous sites in the parental generation and candidate mutations in the F1 or F2 generations that corresponded to known segregating alleles in the known SNPs dataset (e.g. *Sus scrofa* dbSNP dataset) as the reason for the low false-negative rate in the above filter criteria. Similarly, Pfeifer (2017) also found a low false-negative rate after the highly stringent computational filters, particularly for those mutations stably inherited to subsequent generations. Strictly following the mutation-detection procedure of Smeds et al. (2016), we find no reason to expect that the false-negative rate should be significantly different in our study. Given the relatively high depth of coverage, we also consider the rate of false negatives as very low.

In general, high-quality resequencing, stringent bioinformatic pipeline, IGV manual curation and validation via Sanger sequencing ensure the accuracy of DNMs we detected.

### The evolutionary history of Sus scrofa revisited through the de novo mutation rate

We took into account of the results of PSMC and two other methods (MSMC and MSMC-IM), and found some contradictions in previous studies: 1. Two extremely different mutation rates (fig. 4B) were used in the different methods, but the results of divergence time of EUW from ASW were similar; 2. the mutation rates in previous studies led to an incredibly advanced or delayed split of European domesticated pigs from EUW; 3. The Ne of pigs was an order of magnitude less than the other domesticated animals like dogs (Wang et al. 2020), yak (Qiu et al. 2015) and horses (Librado et al. 2015). Additionally, MSMC-IM exhibited ancient evidence of gene flow introgression. Correspondingly, it was reported that at least two events of inter-species admixture occurred when wild boars rapidly spread into Europe, including the migration from Pygmy hogs and a wave of gene flow contributed by an unknown ancient ghost population (Liu et al. 2019). Based on our simulation, we found that the complex gene flow from another Sus species into EUW might uplift the Ne curve and interfere the previous judgment of divergence events based on PSMC. In our case, the date estimation suggested that EUW and ASW actually shared a long history, but the complex migration from an archaic population into EUW mislead us to the deep divergence between EUW and ASW. The phenomenon of introgression influencing the effective population size seems to be common. Hawks (2017) found that the effective population size inferred for particular intervals of time in the past is strongly affected by the history of introgression or gene flow, such as gene flow between non-Africans and Neanderthalian or Denisovan. Inside Africa, introgression also was detected and was suspected from other archaic human groups, with approximately the same inferred divergence date as Neandertals, as suggested by Lachance et al. (2012). Therefore, all modern humans have a clear “wave” of larger inferred effective population size with a “crest”, including Africans (Hawks 2017). A study in dogs also detected imported gene flow lifted the estimated evolutionary trajectory of the target population in PSMC by simulation (Wang et al. 2020). In another recent study, a hump in the historical population size (Ne) estimated with PSMC was attributed to admixture events occurred in donkey (Wang, Li et al. 2020). Hawks (2017) found that the longer the introgression donor diverged from its ancestor, the greater amplitude of the “wave” in the effective population size occurred for the target population. This is consistent with our simulation results. When M = 0.1, the larger Ne rise of target population resulted from introgression from the donor population separates from the ancestral population at 2 Mya (~6 × 10^5^, fig. 5D) than that at 500 Kya (~3 × 10^5^, fig. 5B). The Sus species introgressing into EUW can be traced to at least 1 Myr based on this study (1 - 4 Mya suggested by MSMC-IM, supplementary fig. S5C, Supplementary Material online) and the previous one (Liu et al. 2019). The receptor with archaic admixture from the donor that separates from the ancestral population at 2 Mya in our simulations exhibited a similar trajectory with that of EUW. This probably suggested the origin of introgression donor Sus population can be traced to 2 Mya, and introgression from it into EUW further caused the demographic curves of EUW and SCW separated ~ 2 Mya (supplementary fig. S5A, Supplementary Material online). This independent introgression from archaic population into European wild boars results in a unique phenomenon of PSMC in pigs that the Ne trajectories of SCW and EUW separated significantly before their separated. Therefore, it reminds us to make a demographic inference based on multiple shreds of evidences to avoid the interference of introgression or gene flow.

The de novo mutation rate directly estimated from the pedigree can be used to characterize the demographic history of *Sus scrofa* reliably (Scally and Durbin 2012; Koch et al. 2019). Herein, we used the de novo mutation rate to reconstruct the population history of pigs. Sus speciation occurred on ISEA ~ 1.36 Mya, leading to the emergence of the oldest *Sus scrofa*, then pigs arrived in Eurasia from ISEA and colonized Southeast Asia at ~ 275 Kya. Next, the spread of wild boars into Europe was ~ 219 Kya. The colonization of North China, where the climate was cold, of pigs happened ~ 25 Kya. These estimated population histories were much more recent than the generally accepted history (Frantz et al. 2013), but they were consistent with some evidences, including documentary records about domestication. The new date estimation also could give us new thinking regarding to population genetics of pigs.

The divergence time between European wild and domesticated pig was estimated around 10 Kya using the de novo mutation, which perfectly coincided with the generally accepted domestication time ~10 Kya based on documentary records (Groenen et al. 2012; Frantz et al. 2013; Frantz et al. 2019). We first validated the domestication of pigs ~10Kya by genetic data. The Asian and European wild pigs split 219 Kya (fig. 3B and C; supplementary table S6, Supplementary Material online), far less than the 1.2 Mya (Groenen et al. 2012; Frantz et al. 2013) and suggesting Eurasian wild pigs shared a considerably long same history than we thought before. South (Nanchang, Jiangxi province) and North Chinese wild pigs diverged at 25 Kya, less than 600 Kya (Frantz et al. 2013). Based on this recent split time, we could make a hypothesis that human hunting and domestication activities might accelerate the divergence of North and South Chinese wild pigs and force pigs to migrate into a cold environment, which is a severe challenge to their survival.

The new estimated mutation rate also revealed a maximum effective population size of 2.7 × 10_5_ in pigs, ~6 times larger than that estimated before (Groenen et al. 2012; Frantz et al. 2013) (fig. 3A; supplementary fig. S8, Supplementary Material online) and similar to the population size of other mammals like dogs (Wang et al. 2020), yak (Qiu et al. 2015) and horses (Librado et al. 2015). Our results also revealed a bottleneck in the European wild pigs after colonizing Europe (fig. 3A) rather than a population expansion (Groenen et al. 2012). Similar bottlenecks observed in non-African human populations (Li and Durbin 2011) and Western Eurasian dogs (Frantz et al. 2016) have been interpreted as signs of migration to a new living environment. Penultimate Glacial Period (PGP, 135-194 Kya) followed the western-spread of pigs. Cold climate exacerbated the bottleneck of the western-spreading pigs. Instead, during this period, pigs in Southeast Asia had not yet started to spread northward, and the cold made the pigs gather more in warm areas, which led to a temporary increase of Ne. We have previously found that a 52-Mb segment on X chromosome of European pigs from another genus of pigs may lead to their cold adaptability (Ai et al. 2015). Combined with the results of this study, we speculate that after pigs came out of Southeast Asia, due to the new environment and the following PGP, the western-spreading pigs experienced a great population bottleneck. During this period, the pig hybridized with the other pig genera and got the “gift” to adapt to the cold to a certain extent, then the beneficial introgressed fragment further fixed in the population during their colonization on the European continent. All Sus populations investigated here, even including *Sus cebifrons*, suffered bottlenecks during the Last Glacial Maximum (LGM; 20 Kya; fig. 3A; supplementary fig. S8, Supplementary Material online). The bottleneck of *Sus cebifrons* was not found during LGM in the previous study (Frantz et al. 2013). The bottlenecks observed here were more severe than those reported before (Groenen et al. 2012). Notably, the wild sow is the only ungulate that must build a nest to provide the litter with a warm microenvironment (Algers and Jensen 1990), due to an essential gene, *UCP1*, participating in brown adipose tissue-mediated adaptive nonshivering thermogenesis has been lost ~20 Mya (Berg et al. 2006). The cold climate during the glacial period was fatal to pigs. As expected, we found the lowest Ne of pigs during the LGM.

Altogether, we found some irrationalities and contradictions in the previous estimated evolutionary history of pigs. To address these incompatibilities, we estimated the de novo mutation rate of pigs via a whole-genome three-generation pedigree with nine individuals, which is another non-primate mammalian species obtaining the direct mutation rate after wolves (dogs) and mice. The estimated mutation rate of pigs using a pedigree ultimately didn’t show an abnormally large or small value but one at the same order of magnitude as the mutation rate of other common mammals like wolves (dogs) and yak (fig. 1). This mutation rate enables us to study the population genetics of pigs better than ever before and we re-investigated the population history of pigs with the new mutation rate. Besides, complex ancient admixture could lead to misjudgment of population history, so it is necessary to make a demographic inference based on multiple shreds of evidence. Our results advance the understanding of the population history of pigs.

## Materials and Methods

### Samples and sequencing

We used whole genomes for a known pedigree of nine pigs from a three-generation pedigree (fig. 2; supplementary table S3, Supplementary Material online). Among them, two boars in the parent generation (F0) were White Duroc, sows in the F0 generation were Erhualian. The two boars F0-73 and F0-75 were sequenced (Ai, H. *et al*. manuscript submitted) using the HiSeq 2000 platform. The other seven individuals were sequenced in the same way as the former two pigs. Briefly, genomic DNA was extracted from ear tissues using a standard phenol-chloroform method and then sheared into fragments of 200-800 bp according to the Illumina DNA sample preparation protocol. These treated fragments were end-repaired, A-tailed, ligated to paired-end adaptors and PCR amplified with 500 bp (or 350 bp) inserts for library construction. Sequencing was performed to generate 100 bp (or 150 bp) paired-end reads on a HiSeq 2000 (or 2500) platform (Illumina) according to the manufacturer’s standard protocols. The reads were aligned to the *Sus scrofa* reference genome (build 11.1) using BWA (Li and Durbin 2009) with default options. The mapped reads were subsequently processed by sorting, indel realignment, duplicate marking, and low-quality filtering using Picard (http://picard.sourceforge.net) and GATK 3.5.0 (McKenna et al. 2010; DePristo et al. 2011). To generate an initial trial set of variants for recalibrating base quality scores, we used GATK’s UnifiedGenotyper and SAMtools (Li et al. 2009) to call variant sites, separately, and then took the intersection of variant sites from these two methods. Next, we kept SNP sites if they passed the recommended hard filtering thresholds (QD > 2, FS < 60, MQ > 40, MQRankSum > −12.5, and ReadPosRankSum > 15) as described before (Koch et al. 2019) and filtered sites in repetitive regions from this set using RepeatMasker 4.0.6 (Smit 2013-2015). Finally, we treat the remaining SNPs as a “known” good quality variants set to recalibrate base quality scores using GATK 3.5.0.

After recalibrating base quality scores, the genotypes of all sites were called with GATK’s UnifiedGenotyper with the “emit all sites” options. We did not perform variant quality recalibration (VQSR) suggested by GATK’s best practices since the fact that de novo mutations should only occur in a single individual and are therefore more likely to be filtered out as low-quality variants. Instead, we applied a set of highly stringent hard filter criteria to weed out potential false positives (supplementary table S3, Supplementary Material online). Following Smeds et al. (2016), repetitive regions were masked with a combination of RepeatMasker v3.2.9 (Smit 2013-2015), Tandem Repeats Finder v4.07 (Benson 1999), and a custom Shell script to remove any homopolymers >10 bp that were not already masked (this criterion excluded ~43.4% of the autosomes genome). Then sites passing GATK’s CallableLoci level were kept, and genotype quality (GQ) of each site had to be at least 30 (96.2% - 97.5% of the autosomes genome met this criterion). A hard coverage threshold of 10 was used to minimize false variant calls due to insufficient read data (90.6%–99.1% of the autosomes genome met this criterion) following Keightley et al. (2014). These procedures ensured enough sufficient data to obtain accurate de novo mutations and filter false positives. A total of 1.17-1.28 Gb sequence per individual (51.8% - 56.7% of the autosomes genome) was kept for further quality control of de novo mutations. Finally, we excluded nonvariant sites and indels, only keeping the single nucleotide variants (SNV) in this study (supplementary table S7, Supplementary Material online).

### Identification of de novo mutations

We applied extremely stringent bioinformatic filtering in attempts to have high confidence on the de novo mutations following the previous studies (Kong et al. 2012; Keightley et al. 2014; Smeds et al. 2016; Pfeifer 2017; Koch et al. 2019). Before filtering, we detected a total of 24.3 million SNPs that segregate in the pedigree, concordant with expectations based on previously reported nucleotide levels (Choi et al. 2015). For each individual in the F1 and F2 generations, heterozygous positions were extracted from the background and had to meet the following criteria to be considered as potential de novo mutations:

1. Both parents were required to be homozygous for reference allele with no reads supporting the alternative allele (wipe out the possibility of potential parental mosaicism),
2. No other individuals in the same or the previous generation(s) are heterozygous or homozygous for the alternative allele,
3. At least 25% of the reads support the alternative allele,
4. Does not overlap with the known SNPs from Build 150 of the *Sus scrofa* dbSNP dataset (Sherry et al. 2001) from the NCBI database (Since de novo mutations are rare events, candidates also detected as variation segregating in unrelated individuals are likely false positives).

The filtering criteria in the F2 generation was stricter than that in the F1, which helped reduce more background. The de novo mutations in the F2 generation must simultaneously be homozygous for the reference allele with no reads present that supported the alternative allele in the F0 and the F1 generation. While for the mutation candidates in F1generation, only F0 individuals are required to be homozygous for reference allele with no potential parental mosaicism.

After all these filters, the SNPs left in F1-180, F1-35, F2-1135, F2-1139 and F2-1143 were 393, 360, 180, 213 and 227, respectively.

### Manual curation and annotation of de novo mutations

Candidate mutations were manually curated using the Integrated Genomics Viewer (IGV) (Robinson et al. 2011), similar to Keightley et al. (2014) to visually detect false positives caused by misaligned reads, sequencing errors, insertions, and deletions. The type of false positives detected is excluded as described in the examples shown in the supplemental figures S1 through S4 by Keightley et al. (2014). In the curation process, we applied a two-step approach: firstly, we used a 141 bp-wide window in IGV to screen for the reliable de novo mutations; second, we enlarged the window to 261 bp to further confirm whether the mutation was robustly true based on the status of linked loci on the same read of the candidate mutation, and further to determine whether the mutation origin from the father or the mother (supplementary table S1, Supplementary Material online). Sites were annotated using ANNOVAR v2020Apr28 (Wang et al. 2010) with the annotation of the *Sus scrofa* genome (build 11.1).

### PCR for the detection of the de novo mutations

In this study, we designed a pair of specific primers for each of 44 denovo mutations identified in a family composed of 9 individuals (supplementary table S8, Supplementary Material online). The PCR reaction included 2.5 µL of 10X buffer (Mg2+ plus) (TaKaRa, Japan), 1.5 µL of 25 mM MgCl2, 2.0 µL of 2.5 mM dNTP, 1.0 µL of each primer (10 µM), 0.4 µl of Taq DNA polymerase (5U/µl), 50 ng of the genomic DNA, and the final volume was made up to 25 µl with ddH2O. The mixture was then run in a thermocycler under the following conditions: 94°C for 5 min; 26 cycles of 94°C for 30 s, 68°C (−0.5°C/cycle) for 30 s, 72°C for 45 s; 14 cycles of 94°C for 30 s, 55°C for 30 s, 72°C for 45 s; 72°C for 10 min. The PCR products were sequenced on the 3130XL Genetic Analyzer (Applied Biosystem, USA).

### History inferring methods and models

We used PSMC (Li and Durbin 2011) and MSMC (Schiffels and Durbin 2014) to infer population sizes and split times for Sus populations. The Sus populations are all downloaded from the NCBI SRA database (https://www.ncbi.nlm.nih.gov/sra), including *Sus cebifrons* and *Sus scrofa* (supplementary table S5, Supplementary Material online). Among the data of *Sus cebifrons* in the public database, there are two good-quality individuals, allowing us to estimate the split time. *Sus scrofa* consists of wild pigs (Sumatran wild pigs, South China (Nanchang, Jiangxi Province) wild pigs, North China wild pigs, European (Netherlands) wild pigs) and domesticated pigs (Europe: Large White and Mangalica pigs; China: Hetao, Bamei, Min, Jinhua, Bamaxiang and Wuzhishan). The reads from all the above individuals were aligned to the *Sus scrofa* reference genome (build 11.1) using BWA (Li and Durbin 2009). The subsequent steps, including sorting, indel realignment, deduplication are processed via GATK 3.5.0 (McKenna et al. 2010; DePristo et al. 2011).

For population sizes inferring, PSMC requires diploid consensus sequences. The consensus was generated from the ‘pileup’ command of SAMtools software package (Li et al. 2009). Read depth threshold was set as recommended by PSMC’s manual. Then we used the tool ‘fq2psmcfa’ from the PSMC package to create the input file. We used T_max_= 20, n = 64 (“4+50*1+4+6”) following Groenen et al. (2012). Here we adopted the mutation rate and generation time updated in this study.

Split time estimated by MSMC2 requires two-phased genomes each population. SNPs-calling and low-quality filtering were conducted as previously described (Ai et al. 2015). We phased the samples using SHAPEIT (Delaneau et al. 2013). Besides, there were two masks applied here: one was derived by the tool ‘bamCaller.py’ from the MSMC-tools package; the other one included the sites that were masked using Heng Li’s SNPable mask. Then, MSMC was run on four haplotypes (two from each of the two populations) with the “--skipAmbiguous” argument to skip unphased segments of the genome. The time segments were also set to 64, as in PSMC above. Results were scaled to real time by applying a mutation rate of 3.6 ×10^−9^ per site per generation and a generation time of 3 years derived in this study. MSMC-IM, fitting a continuous Isolation-Migration model to coalescence rates to obtain a time-dependent estimate of gene flow within and across pairs of populations based on the results of MSMC, was also used to decide the time of a split event, presented by a signal of strong gene flow (Wang et al. 2020).

A *split-with-archaic-admixture* model, in which two populations of varying sizes were assumed separated 219 Kya, with one receiving a varying level of gene flow from more deeply diverged population (fig. 5A and C), was built to check how the archaic admixture would affect the shape of PSMC. The ms software (Hudson 2002) was adpoted to perform a series of simulations under the *split-with-archaic-admixture* model. We run the simulations under two different scenes where the third population diverged from the ancestor of the former two populations 500 Kya and 2 Mya, respectively. 10,000 was set as the initial population size used to scale the parameters in the simulation.

## Acknowledgments

This work was financially supported by Innovative Research Team in University (IRT1136), the National Natural Science Foundation of China (31672383), and the National Swine Industry and Technology system of China (nycytx-009). We thank Mingshan Wang from UC, Santa Cruz for helpful comments and suggestions. M.Z. thanks Rasmus Nielsen for useful comments and for hosting him from Januray 2019 to September 2020 at the Center for Theoretical Evolutionary Genomics, UC Berkeley. We also thank for Jiaqi Chen for help on IGV application.

## Author contributions

L.H. organized and coordinated the research. L.H. and H.A. designed the study. M.Z. and H. A. performed the bioinformatics and evolutionary history analyses. Q.Y. was in charge of the DNMs validation by Sanger sequencing. L.H, H.A. and M.Z. analyzed the results. H.A. and M. Z. wrote the draft manuscript. L.H. and H.A. revised the paper.

## References

Ai H, Fang X, Yang B, Huang Z, Chen H, Mao L, Zhang F, Zhang L, Cui L, He W, et al. 2015. Adaptation and possible ancient interspecies introgression in pigs identified by whole-genome sequencing. Nat. Genet. 47(3):217–225.

Algers B, Jensen P. 1990. Thermal microclimate in winter farrowing nests of free-ranging domestic pigs. Livest. Prod. Sci. 25(1-2):177–181.

Benson G. 1999. Tandem repeats finder: a program to analyze DNA sequences. Nucleic Acids Res. 27(2):573–580.

Berg F, Gustafson U, Andersson L. 2006. The uncoupling protein 1 gene (UCP1) is disrupted in the pig lineage: a genetic explanation for poor thermoregulation in piglets. PLoS Genet. 2(8):e129.

Bosse M, Megens HJ, Madsen O, Frantz LA, Paudel Y, Crooijmans RP, Groenen MA. 2014. Untangling the hybrid nature of modern pig genomes: a mosaic derived from biogeographically distinct and highly divergent Sus scrofa populations. Mol. Ecol. 23(16):4089–4102.

Canu A, Scandura M, Merli E, Chirichella R, Bottero E, Chianucci F, Cutini A, Apollonio M. 2015. Reproductive phenology and conception synchrony in a natural wild boar population. Hystrix 26(2).

Choi J-W, Chung W-H, Lee K-T, Cho E-S, Lee S-W, Choi B-H, Lee S-H, Lim W, Lim D, Lee Y-G. 2015. Whole-genome resequencing analyses of five pig breeds, including Korean wild and native, and three European origin breeds. DNA Res. 22(4):259–267.

Comer CE, Mayer JJ. 2009. Wild pig reproductive biology. Wild pigs: biology, damage, control techniques, and management 51–75.

Delaneau O, Zagury JF, Marchini J. 2013. Improved whole-chromosome phasing for disease and population genetic studies. Nat. Methods 10(1):5–6.

DePristo MA, Banks E, Poplin R, Garimella KV, Maguire JR, Hartl C, Philippakis AA, Del Angel G, Rivas MA, Hanna M. 2011. A framework for variation discovery and genotyping using next-generation DNA sequencing data. Nat. Genet. 43(5):491.

Francioli LC, Polak PP, Koren A, Menelaou A, Chun S, Renkens I, Van Duijn CM, Swertz M, Wijmenga C, Van Ommen G. 2015. Genome-wide patterns and properties of de novo mutations in humans. Nat. Genet. 47(7):822–826.

Frantz LA, Haile J, Lin AT, Scheu A, Geörg C, Benecke N, Alexander M, Linderholm A, Mullin VE, Daly KG. 2019. Ancient pigs reveal a near-complete genomic turnover following their introduction to Europe. Proc. Natl. Acad. Sci. USA. 116(35):17231–17238.

Frantz LA, Mullin VE, Pionnier-Capitan M, Lebrasseur O, Ollivier M, Perri A, Linderholm A, Mattiangeli V, Teasdale MD, Dimopoulos EA. 2016. Genomic and archaeological evidence suggest a dual origin of domestic dogs. Science 352(6290):1228–1231.

Frantz LA, Schraiber JG, Madsen O, Megens HJ, Bosse M, Paudel Y, Semiadi G, Meijaard E, Li N, Crooijmans RP. 2013. Genome sequencing reveals fine scale diversification and reticulation history during speciation in Sus. Genome Biol. 14(9):1719–1728.

Freedman AH, Gronau I, Schweizer RM, Ortega-Del Vecchyo D, Han E, Silva PM, Galaverni M, Fan Z, Marx P, Lorente-Galdos B. 2014. Genome sequencing highlights the dynamic early history of dogs. PLoS Genet. 10(1):e1004016.

Galtier N, Rousselle M. 2020. How much does Ne vary among species? bioRxiv 861849.

Giuffra E, Kijas JM, Amarger V, Carlborg O, Jeon JT, Andersson L. 2000. The origin of the domestic pig: independent domestication and subsequent introgression. Genetics 154(4):1785–1791.

Gongora J, Cuddahee RE, Nascimento FFd, Palgrave CJ, Lowden S, Ho SY, Simond D, Damayanti CS, White DJ, Tay WT. 2011. Rethinking the evolution of extant sub-Saharan African suids (Suidae, Artiodactyla). Zool. Scr. 40(4):327–335.

Groenen MA. 2016. A decade of pig genome sequencing: a window on pig domestication and evolution. Genet Sel Evol 4823.

Groenen MA, Archibald AL, Uenishi H, Tuggle CK, Takeuchi Y, Rothschild MF, Rogel-Gaillard C, Park C, Milan D, Megens HJ, et al. 2012. Analyses of pig genomes provide insight into porcine demography and evolution. Nature 491(7424):393–398.

Hawks J. 2017. Introgression makes waves in inferred histories of effective population size. Hum. Biol. 89(1):67–80.

Heptner VG, Nasimovich A, Bannikov AGe, Hoffmann RS. 1988. Mammals of the soviet union: Adams Media.

Hershberg R, Petrov DA. 2010. Evidence that mutation is universally biased towards AT in bacteria. PLoS Genet. 6(9):e1001115.

Huang Y, Zhou L, Zhang J, Liu X, Zhang Y, Cai L, Zhang W, Cui L, Yang J, Ji J. 2020. A large-scale comparison of meat quality and intramuscular fatty acid composition among three Chinese indigenous pig breeds. Meat Sci.108182.

Hudson RR. 2002. Generating samples under a Wright–Fisher neutral model of genetic variation. Bioinformatics 18(2):337–338.

Ikegawa S, Mabuchi A, Ogawa M, Ikeda T. 2002. Allele-specific PCR amplification due to sequence identity between a PCR primer and an amplicon: is direct sequencing so reliable? Hum. Genet. 110(6):606–608.

Jing Y, Flad RK. 2002. Pig domestication in ancient China. Antiquity 76(293):724–732.

Keightley PD, Ness RW, Halligan DL, Haddrill PR. 2014. Estimation of the spontaneous mutation rate per nucleotide site in a Drosophila melanogaster full-sib family. Genetics 196(1):313–320.

Keightley PD, Pinharanda A, Ness RW, Simpson F, Dasmahapatra KK, Mallet J, Davey JW, Jiggins CD. 2015. Estimation of the spontaneous mutation rate in Heliconius melpomene. Mol. Biol. Evol. 32(1):239–243.

Kimura M. 1968. Evolutionary rate at the molecular level. Nature 217(5129):624–626.

Koch EM, Schweizer RM, Schweizer TM, Stahler DR, Smith DW, Wayne RK, Novembre J. 2019. De novo mutation rate estimation in wolves of known pedigree. Mol. Biol. Evol. 36(11):2536–2547.

Kong A, Frigge ML, Masson G, Besenbacher S, Sulem P, Magnusson G, Gudjonsson SA, Sigurdsson A, Jonasdottir A, Jonasdottir A. 2012. Rate of de novo mutations and the importance of father’s age to disease risk. Nature 488(7412):471–475.

Kumar S, Subramanian S. 2002. Mutation rates in mammalian genomes. Proc. Natl. Acad. Sci. USA. 99(2):803–808.

Lachance J, Vernot B, Elbers CC, Ferwerda B, Froment A, Bodo J-M, Lema G, Fu W, Nyambo TB, Rebbeck TR. 2012. Evolutionary history and adaptation from high-coverage whole-genome sequences of diverse African hunter-gatherers. Cell 150(3):457–469.

Larson G, Cucchi T, Fujita M, Matisoo-Smith E, Robins J, Anderson A, Rolett B, Spriggs M, Dolman G, Kim T-H. 2007. Phylogeny and ancient DNA of Sus provides insights into neolithic expansion in Island Southeast Asia and Oceania. Proc. Natl. Acad. Sci. USA. 104(12):4834–4839.

Larson G, Dobney K, Albarella U, Fang M, Matisoosmith E, Robins J, Lowden S, Finlayson H, Brand T, Willerslev E. 2005. Worldwide phylogeography of wild boar reveals multiple centers of pig domestication. Science 307(5715):1618–1621.

Larson G, Liu R, Zhao X, Yuan J, Fuller D, Barton L, Dobney K, Fan Q, Gu Z, Liu XH, et al. 2010. Patterns of East Asian pig domestication, migration, and turnover revealed by modern and ancient DNA. Proc Natl Acad Sci U S A 107(17):7686–7691.

Li H, Durbin R. 2009. Fast and accurate short read alignment with Burrows-Wheeler transform. Bioinformatics 25(14):1754–1760.

Li H, Durbin R. 2011. Inference of human population history from individual whole-genome sequences. Nature 475(7357):493–496.

Li H, Handsaker B, Wysoker A, Fennell T, Ruan J, Homer N, Marth G, Abecasis G, Durbin R. 2009. The sequence alignment/map format and SAMtools. Bioinformatics 25(16):2078–2079.

Li M, Tian S, Jin L, Zhou G, Li Y, Zhang Y, Wang T, Yeung CK, Chen L, Ma J. 2013. Genomic analyses identify distinct patterns of selection in domesticated pigs and Tibetan wild boars. Nat. Genet. 45(12):1431.

Librado P, Der Sarkissian C, Ermini L, Schubert M, Jónsson H, Albrechtsen A, Fumagalli M, Yang MA, Gamba C, Seguin-Orlando A. 2015. Tracking the origins of Yakutian horses and the genetic basis for their fast adaptation to subarctic environments. Proc. Natl. Acad. Sci. USA. 112(50):E6889–E6897.

Liu L, Bosse M, Megens H-J, Frantz LA, Lee Y-L, Irving-Pease EK, Narayan G, Groenen MA, Madsen O. 2019. Genomic analysis on pygmy hog reveals extensive interbreeding during wild boar expansion. Nat. Commun. 10(1):1992.

Lynch M. 2010a. Evolution of the mutation rate. Trends Genet. 26(8):345–352.

Lynch M. 2010b. Rate, molecular spectrum, and consequences of human mutation. Proc. Natl. Acad. Sci. USA. 107(3):961–968.

McKenna A, Hanna M, Banks E, Sivachenko A, Cibulskis K, Kernytsky A, Garimella K, Altshuler D, Gabriel S, Daly M, et al. 2010. The Genome Analysis Toolkit: a MapReduce framework for analyzing next-generation DNA sequencing data. Genome Res. 20(9):1297–1303.

Nachman MW, Crowell SL. 2000. Estimate of the mutation rate per nucleotide in humans. Genetics 156(1):297–304.

Nuijten RJ, Bosse M, Crooijmans RP, Madsen O, Schaftenaar W, Ryder OA, Groenen MA, Megens H-J. 2016. The use of genomics in conservation management of the endangered visayan warty Pig (Sus cebifrons). International journal of genomics 2016.

Pfeifer SP. 2017. Direct estimate of the spontaneous germ line mutation rate in African green monkeys. Evolution 71(12):2858–2870.

Qiu Q, Wang L, Wang K, Yang Y, Ma T, Wang Z, Zhang X, Ni Z, Hou F, Long R. 2015. Yak whole-genome resequencing reveals domestication signatures and prehistoric population expansions. Nat. Commun. 6(1):1–7.

Robinson JT, Thorvaldsdóttir H, Winckler W, Guttman M, Lander ES, Getz G, Mesirov JP. 2011. Integrative genomics viewer. Nat. Biotechnol. 29(1):24–26.

Scally A, Durbin R. 2012. Revising the human mutation rate: implications for understanding human evolution. Nat. Rev. Genet. 13(10):745–753.

Schiffels S, Durbin R. 2014. Inferring human population size and separation history from multiple genome sequences. Nat. Genet. 46(8):919.

Sherry ST, Ward M-H, Kholodov M, Baker J, Phan L, Smigielski EM, Sirotkin K. 2001. dbSNP: the NCBI database of genetic variation. Nucleic Acids Res. 29(1):308–311.

Singer FJ. 1981. Wild pig populations in the national parks. Environmental Management 5(3):263–270.

Smeds L, Qvarnström A, Ellegren H. 2016. Direct estimate of the rate of germline mutation in a bird. Genome Res. 26(9):1211–1218.

Smit A, Hubley, R. & Green, P. 2013-2015. RepeatMasker Open-4.0. <http://www.repeatmasker.org>.

Takashi M, Carl-Johan R, Miguel C, Erik A, Leif A, Webster MT. 2018. Elevated proportions of deleterious genetic variation in domestic animals and plants. Genome Biol. Evol.(1):276–290.

Walters EM, Wells KD, Bryda EC, Schommer S, Prather RS. 2017. Swine models, genomic tools and services to enhance our understanding of human health and diseases. Lab Anim. 46(4):167–172.

Wang C, Li H, Guo Y, Huang J, Sun Y, Min J, Wang J, Fang X, Zhao Z, Wang S. 2020. Donkey genomes provide new insights into domestication and selection for coat color. Nat. Commun. 11(1):1–15.

Wang G-D, Zhai W, Yang H-C, Wang L, Zhong L, Liu Y-H, Fan R-X, Yin T-T, Zhu C-L, Poyarkov AD. 2016. Out of southern East Asia: the natural history of domestic dogs across the world. Cell Res. 26(1):21–33.

Wang H, Zhu X. 2014. De novo mutations discovered in 8 Mexican American families through whole genome sequencing. BMC Proc. 8(S1):S24.

Wang K, Li M, Hakonarson H. 2010. ANNOVAR: functional annotation of genetic variants from high-throughput sequencing data. Nucleic Acids Res. 38(16):e164–e164.

Wang K, Mathieson I, O’Connell J, Schiffels S. 2020. Tracking human population structure through time from whole genome sequences. PLoS Genet. 16(3):e1008552.

Wang MS, Wang S, Li Y, Jhala Y, Thakur M, Otecko NO, Si J-F, Chen H-M, Shapiro B, Nielsen R, et al. 2020. Ancient Hybridization with an Unknown Population Facilitated High-Altitude Adaptation of Canids. Mol. Biol. Evol.

Wang X. 2012. A breeding method of Wannan wild boar in China. In: CN102630633A, editor.

Xiang H, Gao J, Cai D, Luo Y, Yu B, Liu L, Liu R, Zhou H, Chen X, Dun W. 2017. Origin and dispersal of early domestic pigs in northern China. Scientific reports 7(1):1–9.

Yang B, Cui L, Perez-Enciso M, Traspov A, Crooijmans RP, Zinovieva N, Schook LB, Archibald A, Gatphayak K, Knorr C. 2017. Genome-wide SNP data unveils the globalization of domesticated pigs. Genetics Selection Evolution 49(1):71.

