## Supplemental text for "Revisiting the evolutionary history of pigs via de novo mutation rate estimation by deep genome sequencing on a three-generation pedigree"

### Supplementary Figures

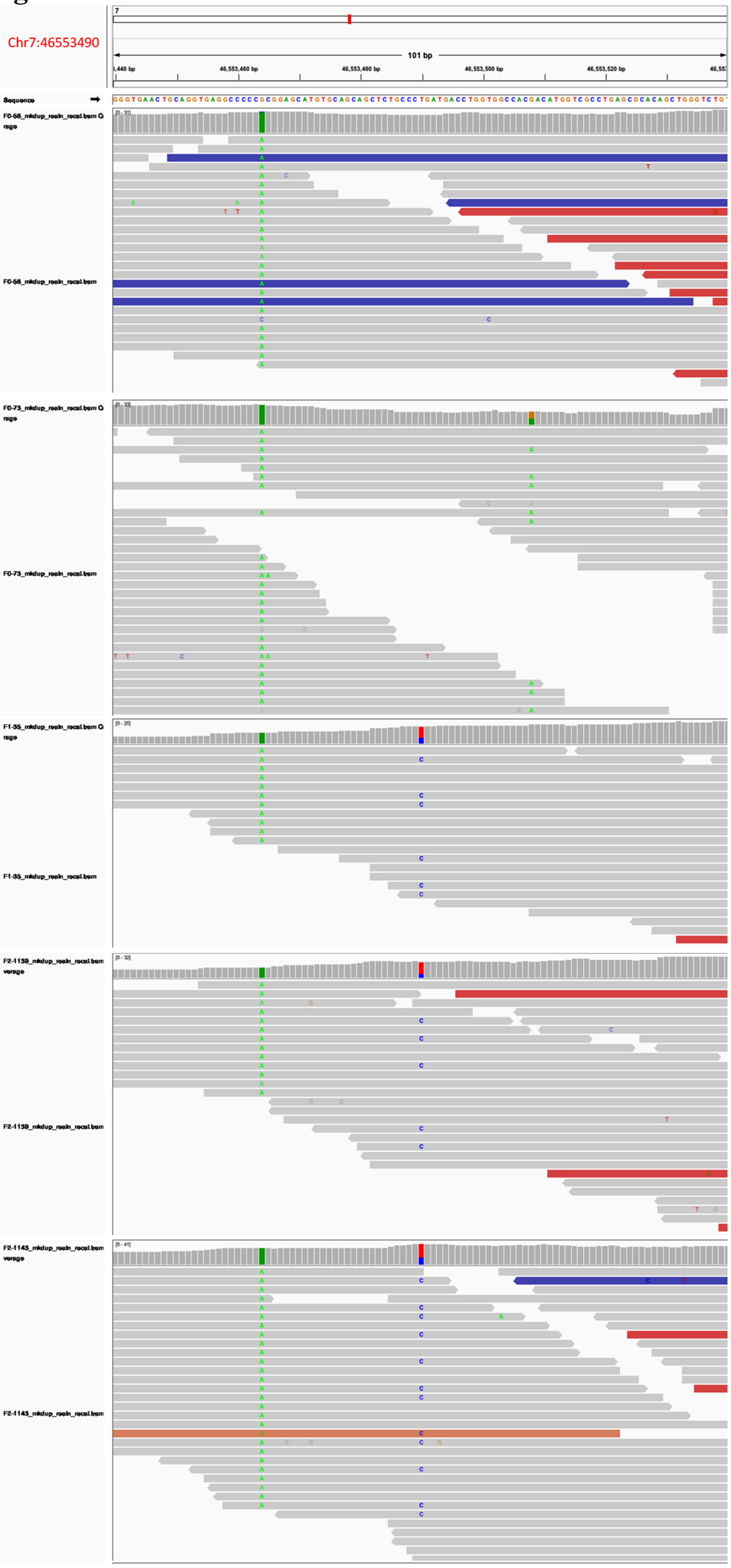


**Figure S1. Screenshot from the IGV showing genotypes at chr7:46553490.**

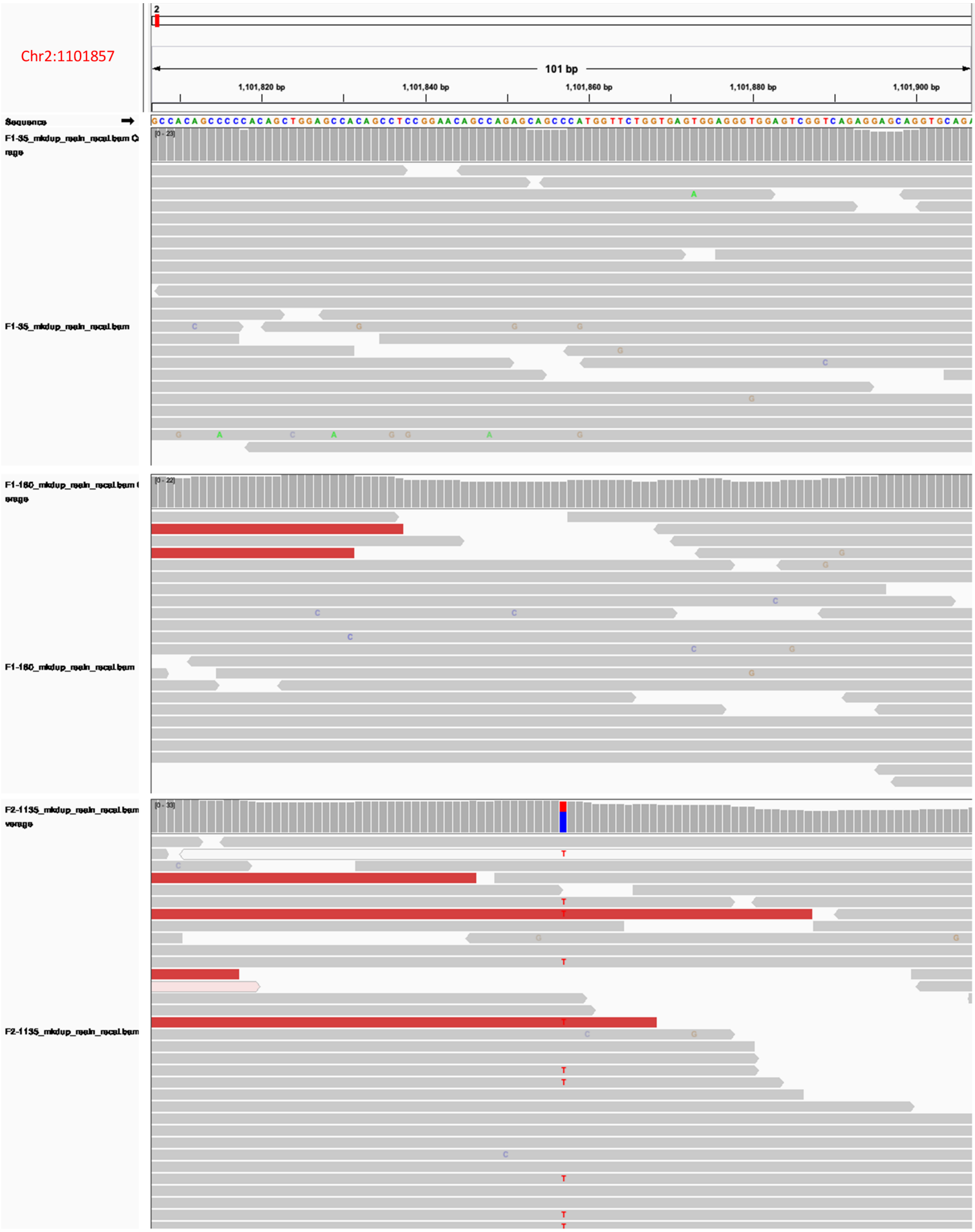


**Figure S2. Screenshot from the IGV showing genotypes at chr2:1101857.**

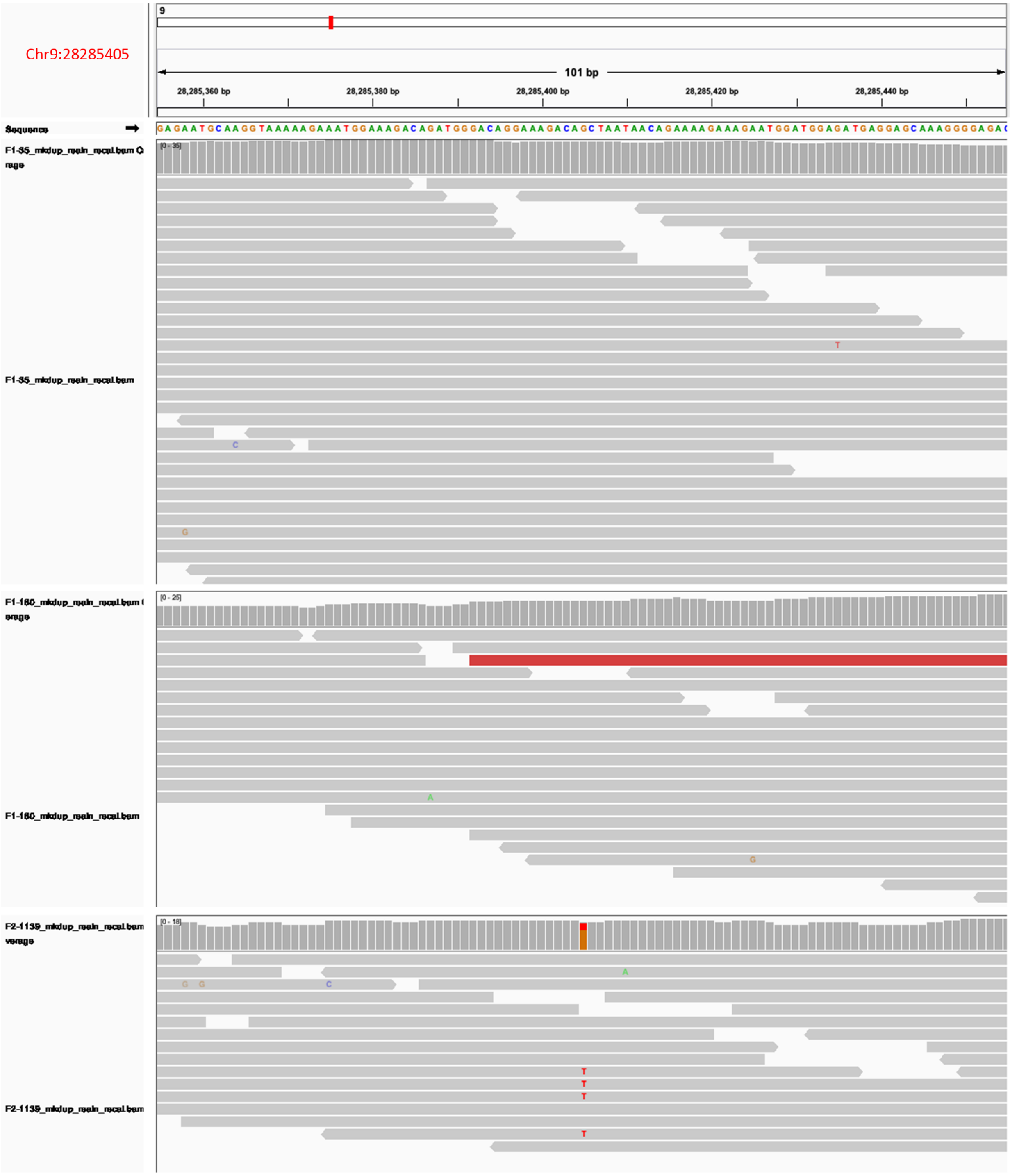


**Figure S3. Screenshot from the IGV showing genotypes at chr9:28285405.**

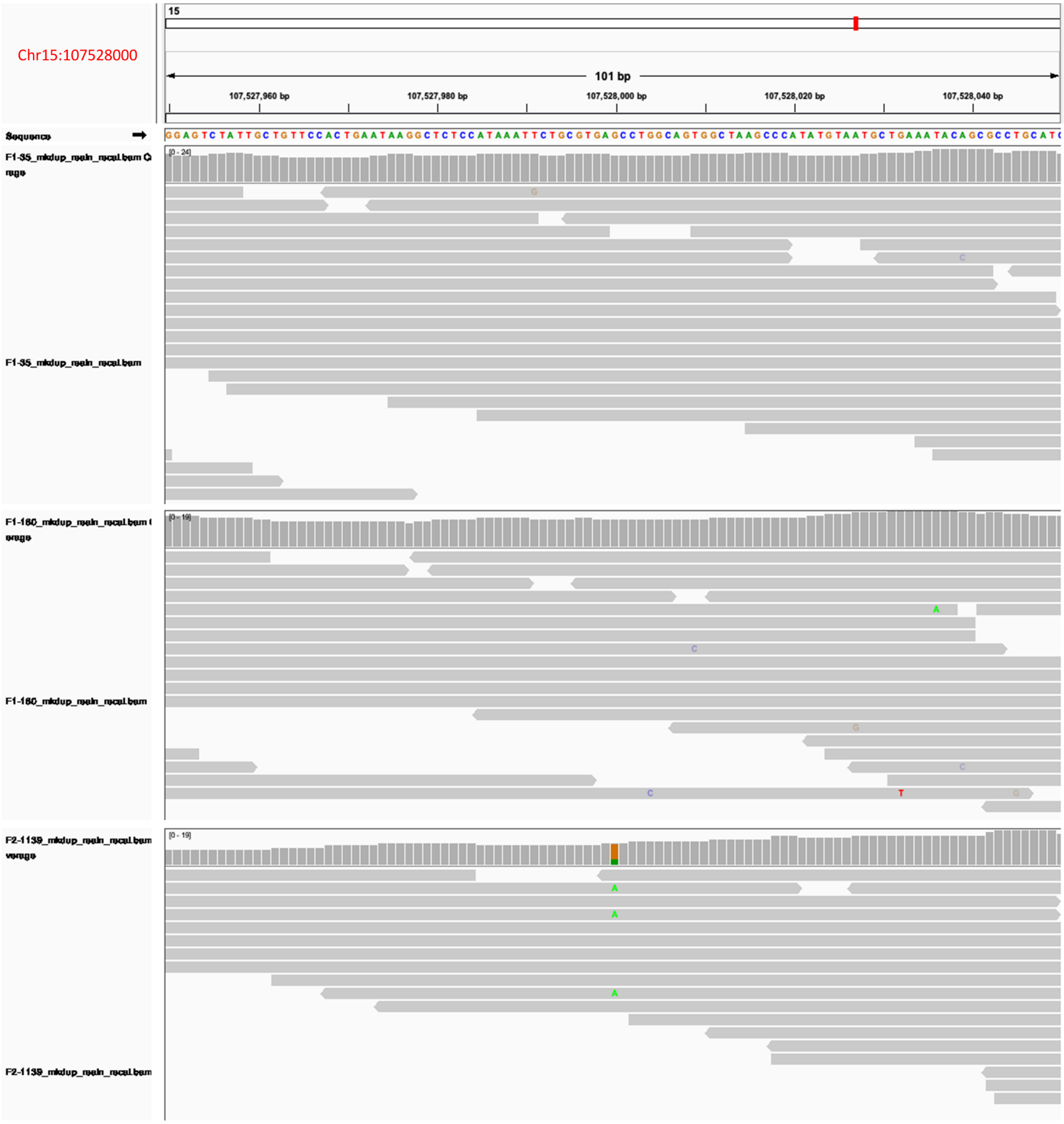


**Figure S4. Screenshot from the IGV showing genotypes at chr15:107528000.**

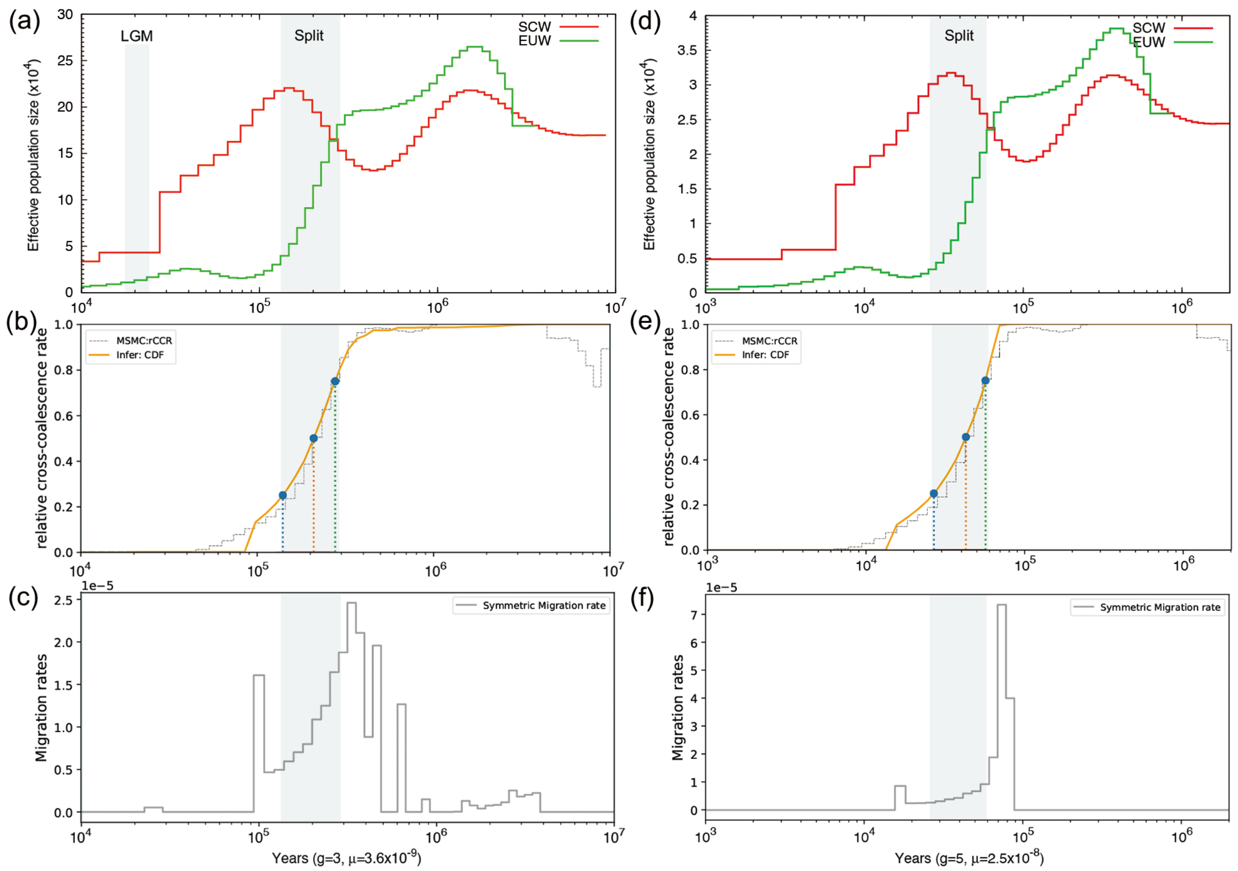


**Figure S5. Comparison of population histories inferred using different mutation rates by PSMC, MSMC and MSMC-IM.** The split time is identified and highlighted in grey at the first cross-over of population size's curves, instead of the second cross-over (backward in time). (a) and (d) Demographic history was inferred with two different mutation rates: $3.6 \times 10^{-9}$ are the mutation rate estimated in this study; $2.5 \times 10^{-8}$ are the mutation rate of human, which was widely used in pig demography inference. The Last Glacial Maximum (LGM) is highlighted in grey in (a). (b) and (e) are inferred by MSMC with the two different mutation rates, respectively. (b) and (e) indicated the split time by the relative cross-coalescence rate (RCCR, orange solid line). The vertical dotted lines represented the date corresponding to RCCR of 25%, 50% and 75%, respectively. (c) and (f) are inferred by MSMC-IM with the two different mutation rates, respectively. (c) and (f) reflected the split time via signals of strong gene flow across SCW and EUW. SCW, South Chinese wild pigs; EUW, European wild pigs.

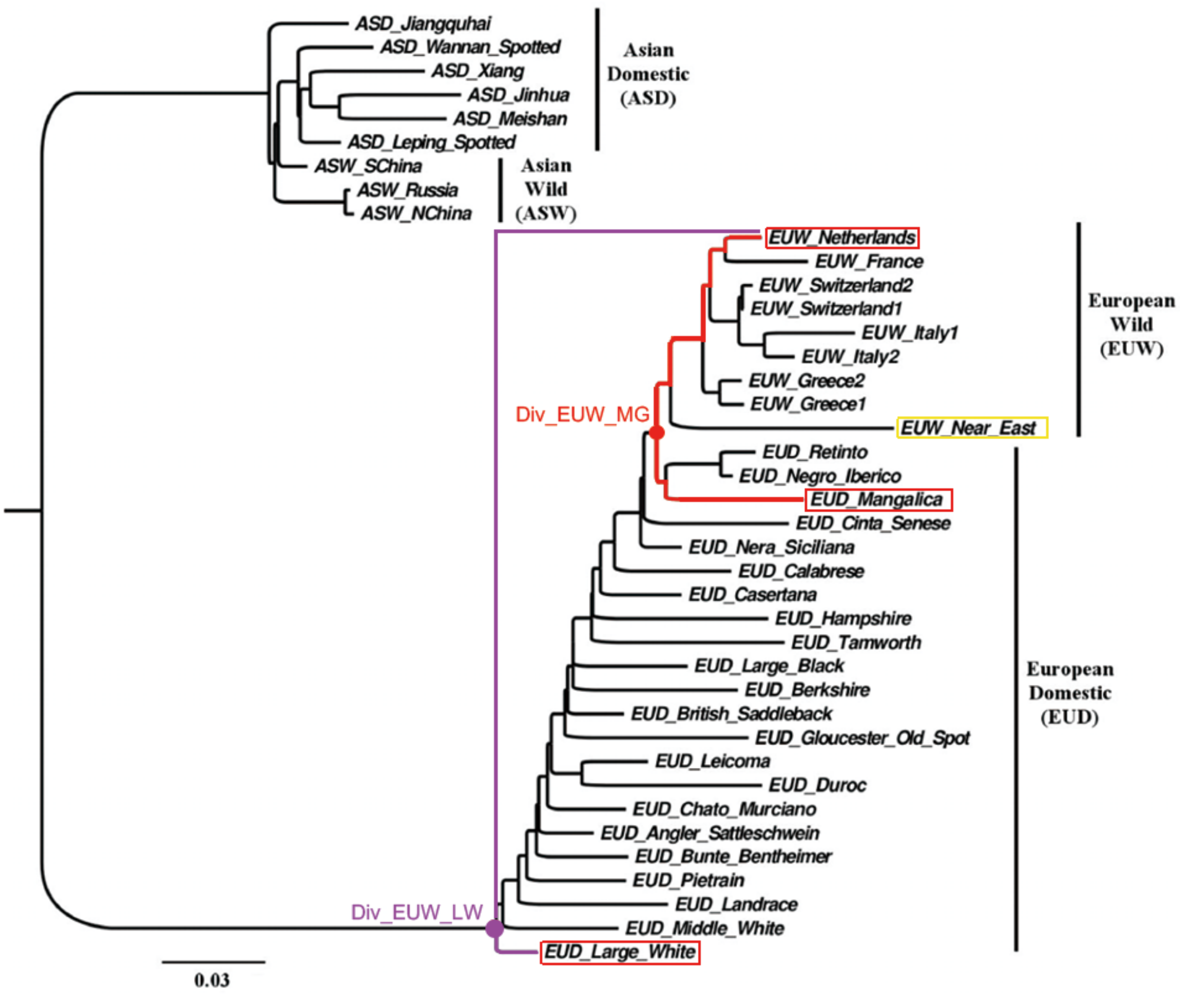


**Figure S6. The pigs sampled for divergence time estimation between European wild and domesticated pigs.** The phylogenetic tree was constructed by TreeMix using data in Frantz, et al. (2015), where the pigs were genotyped by the porcine 60SNP array set. Netherlands wild pigs were selected to be paired with European domesticated pigs, including Large White and Mangalica pigs, to estimate the divergence time. Large White and Mangalica pigs located at the most proximal and distal branch relative to the cluster of European wild pigs, respectively.

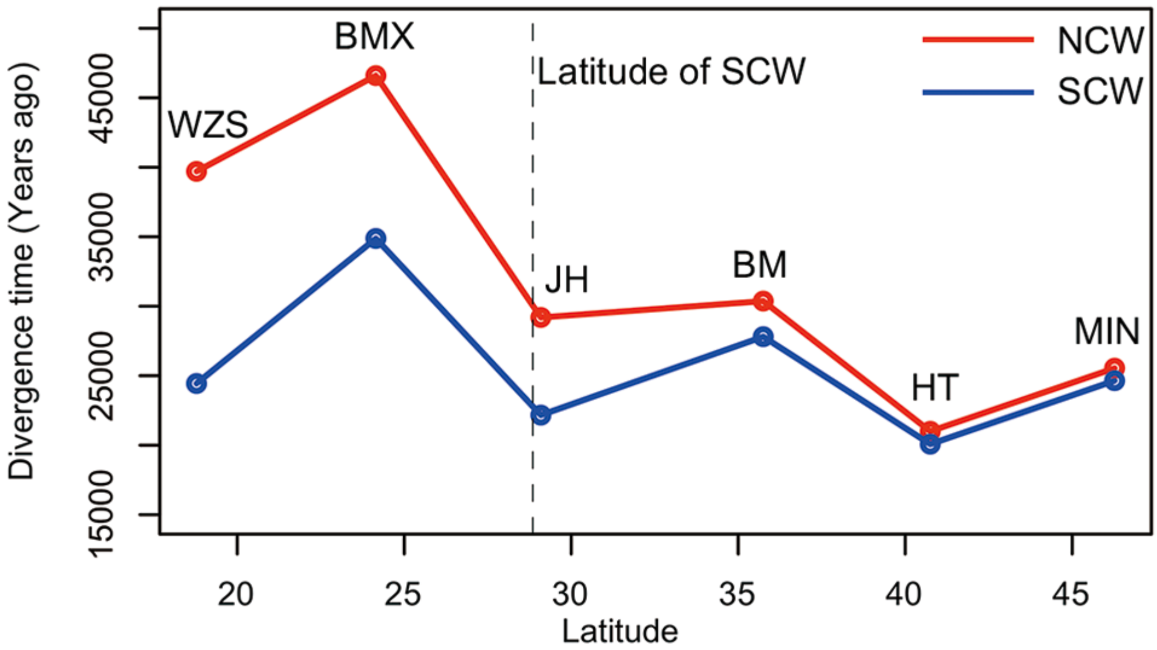


**Figure S7. The relationship between the geographical positions of different Chinese domesticated pig breeds and their split time compared with NCW and SCW, respectively.** The split time here is derived from MSMC. The dotted line represents the latitude of SCW. We didn't know the exact position of NCW (which was used to detect the split time between North and South Chinese pigs in Frantz, et al. (2013), and the only sequenced data available in North China in the public database). The split time of all the Chinese domesticated pigs (including ones distributed in North and South) with SCW are closer than that of NCW. Based on this figure, we thought the place where HT is distributed, beside Yellow River, is a possible place of domestication, echoing the views by Larson, et al. (2010) Thus, we set the split time (20,075 years ago) between HT and SCW as the domesticated time of the Chinese pigs. We also noticed that the time of pigs spreading to North China (25,071 years ago) is close to the domestication time. SCW, South Chinese wild pigs; NCW, North Chinese wild pigs; WZS, Wuzhishan pigs; BMX, Bama Xiang pigs; JH, Jinhua pigs; BM, Bamei pigs; HT, Hetao pigs; MIN, Min pigs (see Supplementary information, Table S5 online, for more detail of per breed).

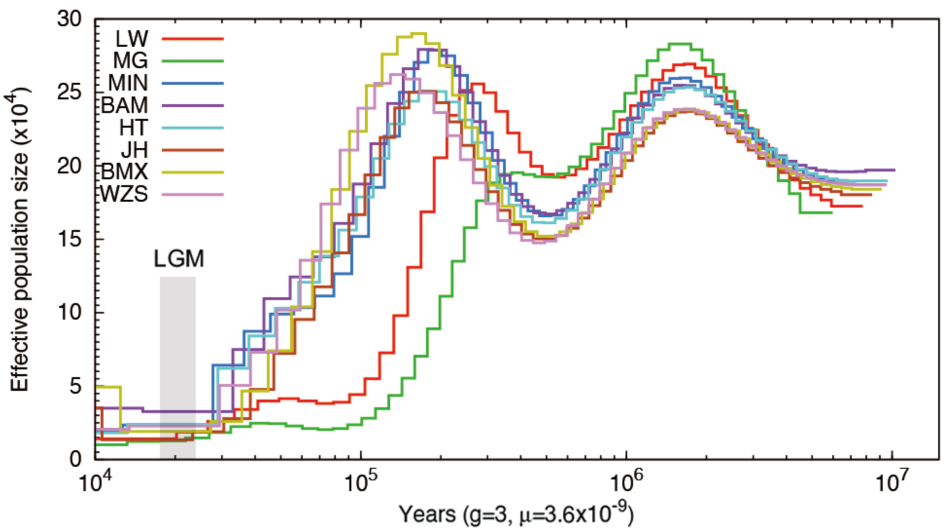


**Figure S8. Dynamic changes of effective population size over past year inferred by PSMC.** The Last Glacial Maximum (LGM) is highlighted in grey. See Supplementary information, Table S5 online, for more detail of per breed. LW, Largewhite pigs; MG, Manglica pigs; WZS, Wuzhishan pigs; BMX, Bama Xiang pigs; JH, Jinhua pigs; BM, Bamei pigs; HT, Hetao pigs; MIN, Min pigs (see Supplementary information, Table S5 online, for more detail of per breed).

### References

Frantz LA, Schraiber JG, Madsen O, Megens HJ, Bosse M, Paudel Y, Semiadi G, Meijaard E, Li N, Crooijmans RP. 2013. Genome sequencing reveals fine scale diversification and reticulation history during speciation in Sus. *Genome Biol.* 14(9):1719-1728.

Frantz LA, Schraiber JG, Madsen O, Megens HJ, Cagan A, Bosse M, Paudel Y, Crooijmans RP, Larson G, Groenen MA. 2015. Evidence of long-term gene flow and selection during domestication from analyses of Eurasian wild and domestic pig genomes. *Nat. Genet.* 47(10):1141-1148.

Larson G, Liu R, Zhao X, Yuan J, Fuller D, Barton L, Dobney K, Fan Q, Gu Z, Liu XH, et al. 2010. Patterns of East Asian pig domestication, migration, and turnover revealed by modern and ancient DNA. *Proc Natl Acad Sci U S A* 107(17):7686-7691.
